# The Cauchy Process on Phylogenies: a Tractable Model for Pulsed Evolution

**DOI:** 10.1101/2023.04.05.535685

**Authors:** Paul Bastide, Gilles Didier

## Abstract

Phylogenetic comparative methods use random processes, such as the Brownian Motion, to model the evolution of continuous traits on phylogenetic trees. Growing evidence for non-gradual evolution motivated the development of complex models, often based on Lévy processes. However, their statistical inference is computationally intensive, and currently relies on approximations, high dimensional sampling, or numerical integration. We consider here the Cauchy Process (CP), a particular pure-jump Lévy process in which the trait increment along each branch follows a centered Cauchy distribution with a dispersion proportional to its length. In this work, we derive an exact algorithm to compute both the joint probability density of the tip trait values of a phylogeny under a CP, and the ancestral trait values and branch increments posterior densities in quadratic time. A simulation study shows that the CP generates patterns in comparative data that are distinct from any Gaussian process, and that Restricted Maximum Likelihood (REML) parameter estimates and root trait reconstruction are unbiased and accurate for trees with 200 tips or less. The CP has only two parameters but is rich enough to capture complex pulsed evolution. It can reconstruct posterior ancestral trait distributions that are multimodal, reflecting the uncertainty associated with the inference of the evolutionary history of a trait from extant taxa only. Applied on empirical datasets taken from the Evolutionary Ecology and Virology literature, the CP suggests nuanced scenarios for the body size evolution of Greater Antilles Lizards and for the geographical spread of the West Nile Virus epidemics in North America, both consistent with previous studies using more complex models. The method is efficiently implemented in C with an R interface in package cauphy, that is open source and freely available online. Cauchy process, Phylogenetic Comparative Methods, Evolutionary jumps, Quantitative traits, Phylogeography

---

A key question in comparative biology is to understand how the phenotypic diversity observed among present day organisms has been shaped by past evolutionary mechanisms. Several hypotheses about the nature and the tempo of phenotypic trait evolution have been proposed. As opposed to gradual evolution, that posits changes at a regular pace over long periods of times, pulsed evolution hypothesizes that occasional “jumps” can occur in the trait value in between long periods of stasis (Simpson, 1944; Gould and Eldredge, 1977). Phylogenetic comparative methods (PCMs) have been used to compare the fit of various models of trait evolution on extant data to support one mode of evolution or the other for a wide range of organisms and traits, such as vertebrate morphology (Landis and Schraiber, 2017), microbial genomic traits (Gao and Wu, 2022a), or virus spatial dispersion (Pybus et al., 2012).

Given a fixed, rooted and time calibrated phylogenetic tree, PCMs usually model the evolution of traits in time as a stochastic process running on the branches of the tree. From the well known Brownian Motion (BM, Felsenstein, 1973, 1985), PCMs have been enriched with many models over the years (see e.g., Harmon, 2019, for a review). In particular, Gaussian processes are versatile tools that can model gradual evolution, including with neutral increments with the BM, stabilizing selection with the Ornstein-Uhlenbeck process (OU, Hansen, 1997), or explosive change with the Early Burst (EB, Harmon et al., 2010). They can be extended to accommodate for evolutionary jumps, with shifts in their defining parameters, either fixed before hand (Butler and King, 2004; O’Meara et al., 2006; Beaulieu et al., 2012; Clavel et al., 2015), or automatically detected (Eastman et al., 2011, 2013; Mahler et al., 2013; Uyeda and Harmon, 2014; Khabbazian et al., 2016; Bastide et al., 2017, 2018). These models enjoy nice algorithmic properties, as their likelihood on a tree can be computed in linear time (Mitov et al., 2020; Bastide et al., 2021), however they require a large number of parameters to represent the shifts, and the search in the parameter space for automatic shift detection can be burdensome and lead to identifiability issues (Bastide et al., 2017).

To avoid this parametrization problem, Lévy processes have been used to directly model pulsed evolution (Landis et al., 2013). In addition to a Brownian diffusion part, Lévy processes have a jump part, which gives the distribution of shift events on the trait. This class of models is very rich, and is appealing as it provides us with a theoretically sound way of describing various modes of possibly pulsed evolution. However, the statistical fit of these models is complex, and relies on the stochastic sampling of high-dimensional latent spaces (Bokma, 2008; Lemey et al., 2010; Landis et al., 2013; Elliot and Mooers, 2014; Duchen et al., 2017), and often requires additional numerical integral evaluations. Although recent progresses have been made, using either likelihood approximations (Landis and Schraiber, 2017) or efficient Hamiltonian Monte Carlo sampling (Fisher et al., 2021), these models remain computationally intensive.

As pointed out by Blomberg et al. (2020), the enduring popularity of the simple BM and OU models, despite growing evidence for wide-spread non-gradual evolution (Landis and Schraiber, 2017; Gao and Wu, 2022a), is rooted in their simplicity and tractability. In order to model pulsed evolution, we consider here the Cauchy process (CP) on trees, that is a simple and (now) tractable pure-jump process in which the trait increment along each branch follows an independent centered Cauchy distribution of dispersion proportional to its length.

The Cauchy process is comparable to the Brownian motion in many aspects. Both have only two parameters, namely the root trait value and the process variance (for the BM) or dispersion (for the CP). In addition, both are limiting cases of a Lévy-stable process: where the BM can be seen as a purely gradual model, the CP is a pure-jump process, with no diffusion component, and can be seen as a purely pulsed model. Both models are hence useful to have in our analysis toolbox, and the relative fit of one model against the other can quickly give us an idea of the general pattern of trait evolution on a specific dataset, either gradual or pulsed.

We show here that the CP is tractable, and we provide an explicit recursive algorithm to compute the likelihood of a univariate vector of traits at the tip of a dated and possibly non ultrametric phylogenetic tree, in a single traversal of the tree. We also derive an algorithm to compute the posterior ancestral density, conditionally on tip values, of the trait at internal nodes, and of the increments at all branches (the increment of a branch being the difference between the traits of the child and parent nodes of the branch). To our knowledge, the CP is the first non-Gaussian model that admits such explicit computations. The proposed method is exact in theory, and we show that it works well for empirical as well as simulated trees with 200 tips or less. However, because of its complex structure, our algorithm suffers from numerical robustness issues, and we empirically observed that it could give unreliable results for larger trees. We provide here some guidelines and diagnostic tools to asses whether computations succeeded on a particular tree.

An important feature of the CP is that, contrary to Gaussian models that always give rise to Gaussian posterior ancestral state distributions (Royer-Carenzi and Didier, 2016), it can reconstruct multimodal trait distributions at ancestral nodes (such a behavior was observed for other Lévy processes in Elliot and Mooers 2014). The different reconstructed modes are well suited to represent the various evolutionary scenarios that are compatible with the observed dataset on extant species, and are an elegant way to handle the identifiability issues that arise in shifted Gaussian models (Bastide et al., 2017).

The article is organized as follows. We first outline the mathematical and modeling properties of the CP, and present the algorithms for likelihood computation and ancestral state reconstruction. We then assess the quality of the statistical inference using simulations, and exhibit some of the properties of the Cauchy process model with the reanalysis of two datasets, chosen in the fields of evolutionary ecology and virology. We discuss some of the strengths and limitations of our approach in the last section.

## 1 Model

In this section, after recalling basic facts about Gaussian processes, we present three possible characterizations of the CP, then discuss the multimodality of posterior ancestral distribution under the CP. Our description is rooted in the fact that both processes fall in the class of *α-stable* processes, with *α* = 2 for the BM, and *α* = 1 for the CP (see e.g., Klenke, 2014, Chap. 16 for an introduction on stable processes).

### 1.1 Cauchy Process on a Tree

#### 1.1.1 Gaussian Models as Stable Processes

Gaussian phylogenetic models can be characterised in two complementary ways (see e.g., Felsenstein, 2004, Chap. 23-25). Conditionally on a time calibrated phylogenetic tree *𝒯*, they can be first described as a stochastic process, such as the BM, running on the branches of the tree, and splitting into independent processes at branching events. An other way is to consider the heritability distribution on the tree, that is, independently for each branch of the tree, the distribution of *Z*_*n*_ the child node’s trait value conditional on 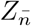 the parent node’s trait value. For a BM with uniform variance *σ*^2^, this conditional distribution is simply Gaussian, centered in the parent’s value, and with a variance that is proportional to the branch length *t*_*i*_, i.e.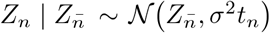. Such heritability formulas are available for more sophisticated Gaussian processes such as the OU or the Brownian with varying rates, and have been used for efficient likelihood and gradient computations (Mitov et al., 2020; Bastide et al., 2021). The equivalency of these two descriptions stems from the fact that the Gaussian distribution is a *stable distribution*, which mean that any linear combination of two independent Gaussian variables is itself a Gaussian variable, with expectation and variance parameters themselves found as linear combinations of the original parameters (Klenke, 2014).

#### 1.1.2 CP as a Stable Process

As the Cauchy distribution is also stable, the most natural way to describe the CP is to use the branch heritability description, simply replacing the Gaussian by the Cauchy distribution:

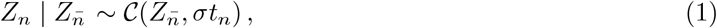

in other words, the distribution of the child node’s trait value is Cauchy, with location parameter the parent node’s trait value, and a scale parameter proportional to the length of the branch. The CP density is explicit (see Equation (2)), and, similarly to the BM, it has two parameters: *μ* the trait value at the root of the tree, and *σ* the dispersion of the distribution. Under the CP, conditionally on the root trait value and at point of the tree, the trait follows a Cauchy distribution with dispersion proportional to its distance from the root, instead of a Gaussian distribution. The Cauchy distribution does not have any finite expectation or variance. This implies that the CP, no matter the timescale or the parameter values, will never converge to a Gaussian distribution, as opposed to other Lévy processes (Landis et al., 2013). As shown in the discussion, the CP is however itself a natural limit for several other stochastic processes. As the Cauchy distribution is heavy-tailed, the child trait value can be quite different from its parent value, which can be associated to jumps in the underlying stochastic process (see Fig. 1).

**Figure 1:**
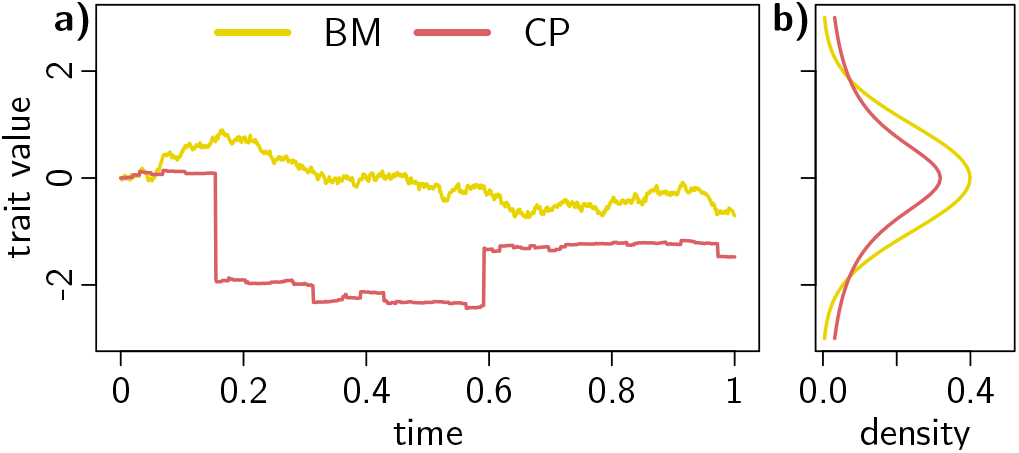
The Cauchy process (CP, dark red) is a pure jump process and produces a trait distribution with fatter tails, compared to the Brownian Motion (BM, light yellow) that is continuous. **(a)** Realization of a CP and BM with initial value 0 and dispersion or variance parameter 1 on a unit time interval. (b) Associated distribution of the trait at the end of the interval, either a standard Cauchy or standard Normal.

#### 1.1.3 CP as a Pure Jump Process

Indeed, as mentioned before, this model is associated to a stable random process on the tree. This process is a *pure jump* Lévy process, that can be described as an infinite sum of independent compound Poisson processes, each with jump rate of 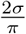, and jump distribution 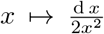 on an interval 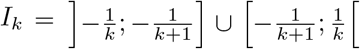, for *k ≥* 0, with the convention that 1*/*0 = *∞*. More formally, the Lévy-Khinchine representation is given by the triplet (0, 0, *v*), with 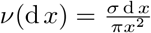 (Klenke, 2014). The CP has hence null diffusion and drift components, but constantly jumps according to distribution *v*. This means in particular that jumps happen on all along every edge of the tree.

#### 1.1.4 CP as a Relaxed Random Walk

A third description of the CP is to see it as a relaxed BM on the tree, where each branch, ending at a given node *n*, has its own independent random variance scale parameter *ϕ*_*n*_ which follows an inversegamma distribution with shape 1*/*2 and scale *t*_*n*_*/*2, i.e., *ϕ*_*n*_ *∼* Inv-Gamma(1*/*2, *t*(*n/*2), such)that the trait propagation on the tree is Gaussian with a scaled variance: 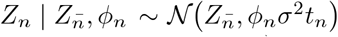. Using the classical normal-inverse-gamma conjugate distribution (see e.g., Gelman et al., 2013), we get that the marginal heredity rule 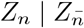 (integrated on the *ϕ*_*n*_) is a Student distribution with one degree of freedom, that is, a Cauchy distribution as in Equation (1).

Note that, in this construction, the distribution of scale parameters *ϕ*_*n*_ depends on the branch lengths *t*_*n*_, which is different from the well-known relaxed random walk (RRW Lemey et al., 2010), that uses independent identically distributed (i.i.d.) *ϕ*_*n*_. In fact, what is usually referred to as the “Cauchy RRW” (Lemey et al., 2010; Pybus et al., 2012), a model that has been widely used in the field of continuous phylogeography (see Baele et al., 2017, for a review), is distinct from the CP as defined here, as it takes *ϕ*_*n*_ *∼* Inv-Gamma(1*/*2, 1*/*2) i.i.d., and leads to a marginal heredity distribution 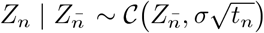, that does not match any Lévy process. In other words, the “Cauchy RRW” amounts to a CP running on the tree obtained by replacing the initial branch lengths by their square root.

### 1.2 Jump Process and Multi-modality of Reconstructed States

Using the CP model, we show in the next section how we can efficiently compute the posterior density conditionally on the trait values at the tips of the tree of both the trait value at internal nodes and the increment magnitude on tree branches. Under non Gaussian stable models such as the CP, Elliot and Mooers (2014) reported several cases of multimodality for this posterior density. We do also observe cases of multimodality of the posterior distribution of both ancestral states and increments under the CP (see Section Empirical Examples). We argue that this phenomenon is strongly connected with the fact that the CP is a jump process, and that the exact position of the large jumps on the tree cannot be determined with certainty from the observed tip values.

In particular, Bastide et al. (2017) showed that under the shifted Brownian process, which is a BM evolutionary model where instantaneous shifts (i.e., jumps) are allowed, several configurations of jumps may lead to the exact same likelihood, and thus are not identifiable under this model. Figure 2-a,b,c displays three jump configurations leading to the same joint distribution of tip values under the shifted BM (a shift changes the expectation of the BM but not its variance). Let us remark that some ancestral nodes do not have the same history in the different scenarios, such as the node marked with a dot on Figure 2, which in scenario (a) has a trait value with expectation 0, the starting value at the root, but has shifted expectations at *−* 1 and 2 in scenarios (b) and (c), respectively.

**Figure 2:**
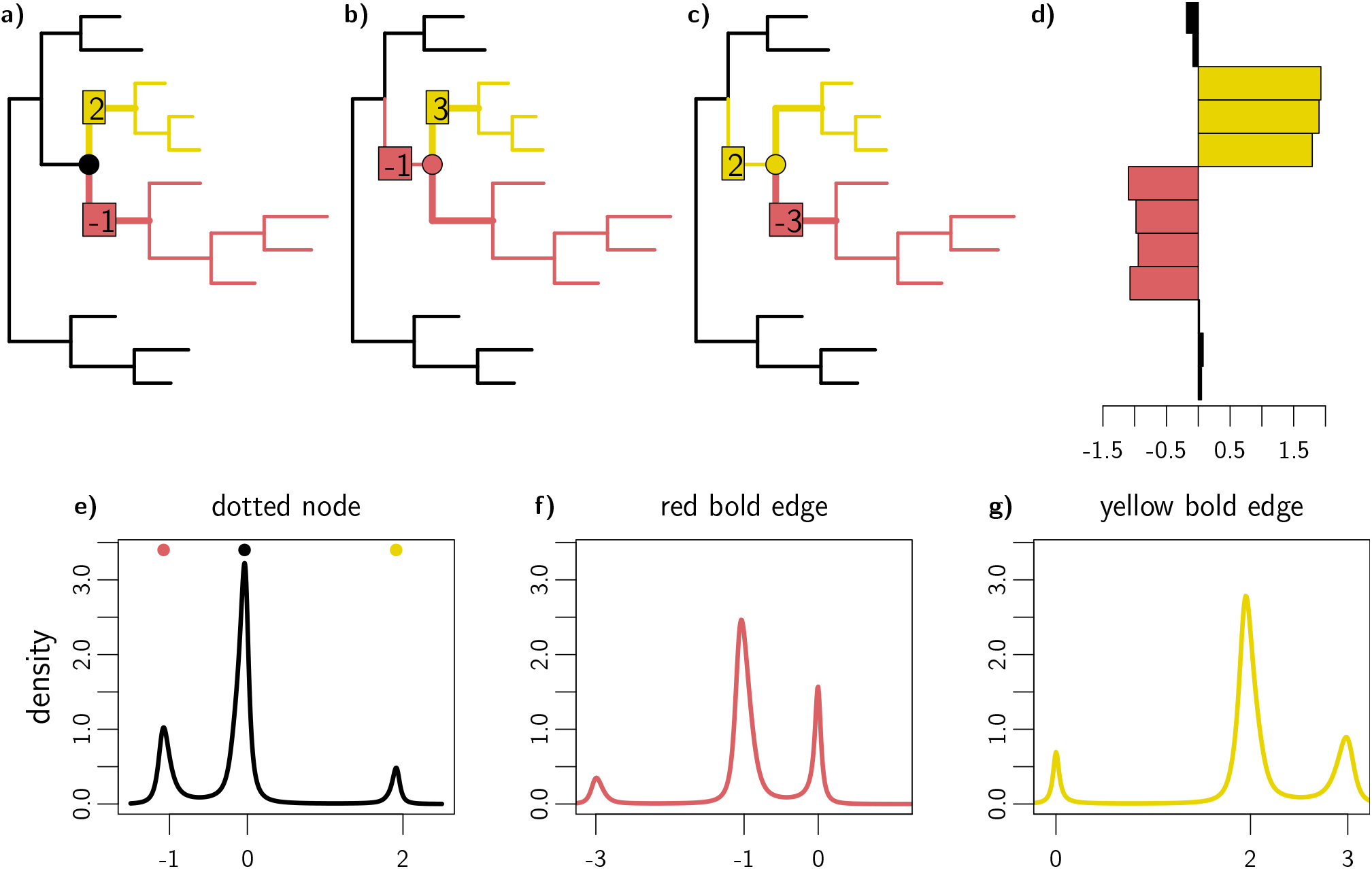
Illustration of the multi-modal ancestral density on a toy example, where the data was simulated according to a simple Brownian Motion on a tree, with shifts on two branches. **a-c)** Three equivalent shift allocations which result in the same trait distribution at the tips. The BM starts with value 0, and then shifts on the colored branches, by the amount showed on the boxes. In all three scenarios, the black tips have expectation 0, the dark red tips expectation -1, and the light yellow tips expectation 2. **d)** Realization of the shifted BM trait distribution at the tips of the tree, with tips colored according to their expectation. **e)** Ancestral reconstruction of the node marked with a colored dot on the trees (a-c), using a Cauchy process fitted on the realization (d). The posterior distribution is tri-modal, with modes at 0 (black), -1 (dark red) or 2 (light yellow), that match the ancestral node expectation under the BM used to simulate the data in the three shift allocations. **f)** Increment reconstruction for the bold dark red branch in trees (a-c), using the same Cauchy process. The posterior distribution is also tri-modal, with modes -1, 0 and -3, matching the shift values on this branch in the three shift allocations. **g)** Increment reconstruction for the bold light yellow branch in trees (a-c). The posterior distribution has modes 2, 3 and 0, matching the shift values on this branch in the three shift allocations.

To illustrate the behavior of the CP in such a situation, we simulated a dataset using one of the shifted BM models presented on Figure 2, and then performed ancestral and increment reconstructions using our CP framework. We observe that these reconstructions reflect all three scenarios (a), (b) and (c). In particular, the posterior density of the “dot-node” state is multimodal, with three modes around to 0, *−* 1 and 2 (Fig. 2-e), which are indeed the ancestral expectations of this node in scenarios (a), (b) and (c), respectively. Similarly, the modes of the posterior increment density on the two daughter branches are consistent with the possible shift values on these branches (Fig. 2-f,g).

As we will see in our empirical datasets analyses below, the CP reconstruction hence provides us with at least two important pieces of information. First, the multimodality of posterior distributions suggests that (significantly) distinct scenarios of trait evolution may have lead to observed trait pattern among extant species. Second, the relative weight of each mode gives us a sense of the uncertainty associated with the different candidate scenarios, an information which was not available under the shifted Brownian model. In particular, if one is only interested in reconstructing a single ancestral value, a natural choice is to consider the highest mode of the posterior distribution.

## 2 Methods

### 2.1 Computation of the Likelihood

#### 2.1.1 Notations and assumptions

Unless stated otherwise, all the trees considered below are dated, possibly non ultrametric, and are *stem trees*, in the sense that they include an extra stem branch ending at their root node. The length of this branch may be zero, in which case the tree is a *crown tree*. For any phylogenetic tree 𝒯, we write *r* for the root node of 𝒯 and L_𝒯_ for its set of tips, with cardinal |L_𝒯_ |. For all nodes *n* of 𝒯, we put 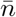 for the direct ancestor of *n* (if it exists) and 𝒯 _*n*_ for the subtree of *𝒯* originating at *n*. Note that the subtree 𝒯 _*n*_ includes the branch ending with *n* in *𝒯* and is thus a stem tree. Without loss of generality, we assume that the tree is binary, allowing for branches of lengths zero to accommodate for possible polytomies. We further assume that a continuous trait is measured as *y*_*a*_ at each tip *a* of 𝒯.

Under the Cauchy model with parameters (*μ, σ*), the evolution process starts with the value *μ* at the beginning of the branch ending with the root of the phylogenetic tree. All the increments Δ_*n*_ along the branches are independent and follow the centered Cauchy distribution with dispersion *σt*_*n*_ where *t*_*n*_ is the length of the branch ending at node *n*, namely with density:

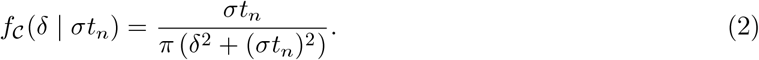

#### 2.1.2 Likelihood Computation

Our main methodological result is to derive a pruning algorithm to compute the joint likelihood of the vector of tip traits values **y** under a CP on the tree.

##### Proposition 1.

*Under the CP with parameters* (*μ, σ*), *the joint likelihood of the vector of trait values* 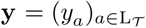 *at the tips of the tree 𝒯 is*

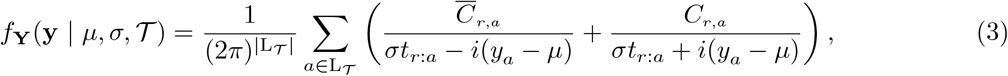

*Where* 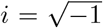 *is the imaginary unit*, 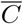 *is the complex conjugate of C, and, for any node n of* 𝒯 *and any tips a of* 𝒯 _*n*_, *t*_*n*:*a*_ *is the sum of the lengths of the branches ending at each node in the path going from n to a on the tree (the length of the branch ending with n included) and the coefficient C*_*n,a*_ *is a complex number recursively computed on 𝒯 as:*

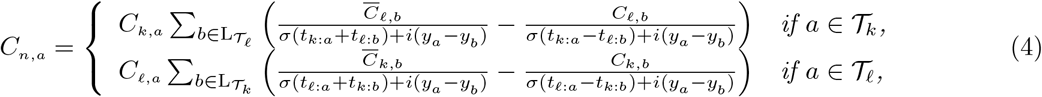

*if n is an internal node and k and l are its two direct descendants, and initializing C*_*a,a*_ = 1 *on all tips a*.

The proof of this algorithm is detailed in the Appendix. It relies on the propagation of partial likelihoods in a postorder on the tree, that takes advantage of the conditional independence structure given by the tree, and exploits closed form integral formulas for the branch integrations. The propagation formulas (4) involve the computation of 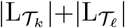 sums at each integration, which leads to a time complexity that is quadratic in the number of tips (see Supplementary Material Section SI-1), and not linear as for Gaussian processes (Mitov et al., 2020).

In our algorithm, the recursion step associated to a node *n* requires to compute a coefficient for all the tips *a* descending from *n*. This computation involves coefficients associated with the tips of the direct subtree of *n* which does not contain *a*. This is quite different from usual recursion formulas in pruning algorithms, such as the one of Felsenstein (1973), that do not imply such cross-computations on separate sub-trees.

### 2.2 Ancestral States Reconstructions

#### 2.2.1 Ancestral Node Reconstruction

Under a Gaussian process, the ancestral trait value at an internal node conditional on all tip values is also Gaussian, so that ancestral state reconstruction reduces to the computation of the conditional expectations and variances of all nodes, which can be done in one single pass on the tree (see e.g., Royer-Carenzi and Didier, 2016). In the Cauchy case, the ancestral distribution cannot be reduced to its moments. However, the Markov property allows us to write the conditional probability density function of the trait *Z*_*n*_ at any internal node *n* as a product of likelihood functions on well-chosen sub-trees:

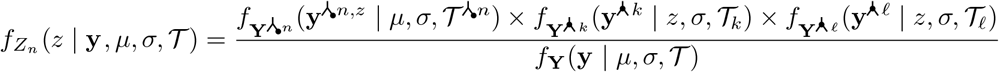

where 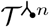 is the tree trimmed after node *n* (Fig. 3-c), with trait values 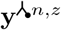 such that 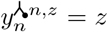 and 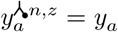 for all tips *a ≠ n* of 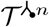, and 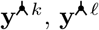 are the vectors of trait values at the tips of sub-trees 𝒯 _*k*_ and 𝒯 _*ℓ*_, with *k* and *ℓ* the two direct descendants of *n*.

**Figure 3:**
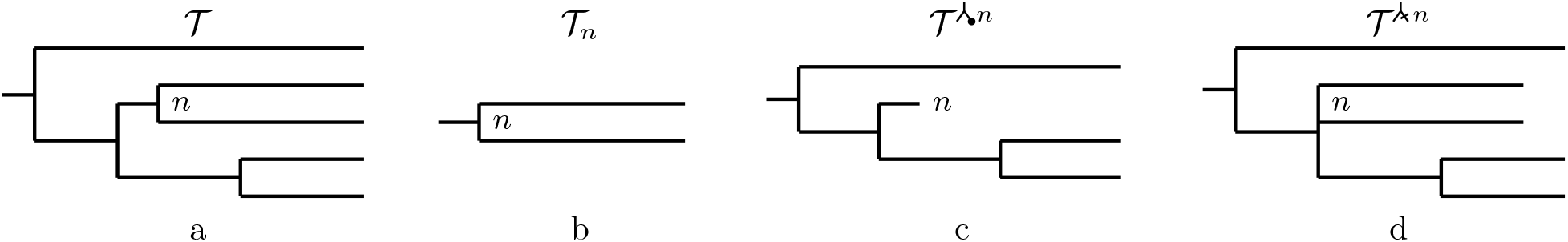
(a) a tree *𝒯* ; (b) the tree *𝒯*_*n*_, i.e., the (stem) subtree of *𝒯* rooted at *n*; (c) the tree 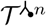, i.e., the tree *𝒯* trimmed at node *n*; (d) the tree 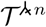, i.e., the tree obtained from *𝒯* by setting the length of the branch ending by *n* to 0.

#### 2.2.2 Branch Increment Reconstruction

Besides ancestral node values, the distribution of the increment Δ_*n*_ of the trait on a branch ending at a given node *n* can help us understand the dynamics of trait evolution. To compute the probability density function of such an increment, we can use a similar decomposition, that relies on canceling out a branch and shifting descending tip values. More precisely, we get:

- for any internal node *n* different from the root, 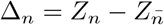, and:

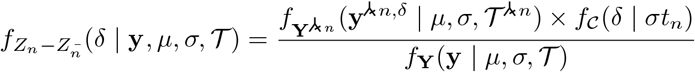

where 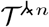 is the tree obtained from 𝒯 by setting the length of the branch ending with *n* to 0 (Fig. 3-d), with tip trait values 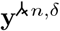 such that 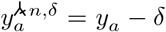 if *a* is a tip of 𝒯 _*n*_, and 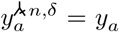 otherwise;
- for the branch ending at the root *r*, if *r* is not a tip, Δ_*r*_ = *Z*_*r*_ *− μ*, and:

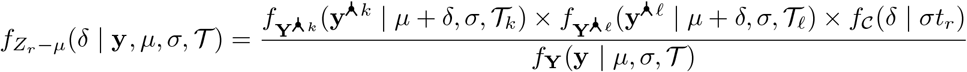

where *k* and *ℓ* are the two direct descendants of *r* and, 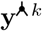 and 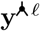 are the vector of tip values of subtrees 𝒯 _*k*_ and 𝒯 _*ℓ*_;:
- in the special case of a tip *a* = *r* or *ā* = *r* and the root branch has length zero, then 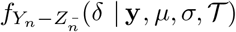 is the Dirac delta function at *y*_*a*_ *− μ*;
- for any other tip *a*, 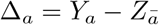, and:

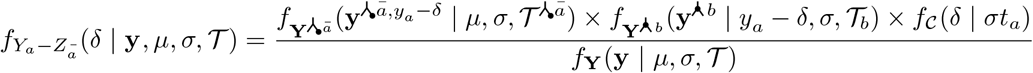

where 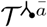 is the tree trimmed after ā the parent of *a*, with tip values 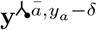 such that 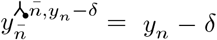 and 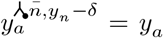 for all tips 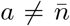 of 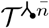, and *b* is the sibling node of *a* with 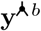 the vector of values at the tips of subtree 𝒯 _*b*_.

Note that the approaches presented in this section are valid and could be used for ancestral state or increment reconstruction for any evolution model satisfying the Markov property.

#### 2.2.3 Computation and Complexity

For each given trait or increment value, the posterior node or branch densities can be computed using the pruning algorithm presented above. In order to visualize them (such as in Fig. 2-e,f,g), one has to perform these computations at many values, which are typically taken from a grid. Supplementary Material Sections SI-2 and SI-3 show how to optimize computations in this situation. In particular, each new evaluation at an internal node or branch is only linear in the number of tips, making these computations efficient even for refined grids.

### 2.3 Stem Trees, Crown Trees and Restricted Maximum Likelihood

In phylogenetic models, the status of the root trait *Z*_*r*_, fixed or random, can have a strong impact on the quality of the statistical inference (see e.g., Felsenstein (1973), and Ho and Ané (2014b) for a discussion in the OU case). Using a stem tree amounts to taking a prior on the root trait, centered on *μ*, and with a variance or dispersion proportional to the stem branch length. This is the default behavior in popular software such as BEAST (Lemey et al., 2010; Suchard et al., 2018), that generally use vague priors on the root, that is, long stem branches, to weaken the dependency to the root. If on the contrary the stem branch has length zero, that is, the tree is a crown tree, then the root is fixed, and the analysis is carried conditionally on the root trait value. Phylogenetic regression frameworks generally use this assumption, and the fixed effects can be formally seen as representing the root trait value (see e.g. R package phylolm, Ho and Ané, 2014a). This root value can however be seen as a nuisance parameter, and, drawing from the linear mixed model literature, the *restricted maximum likelihood* (REML) can be used to marginalize over the root value (see e.g., Searle et al., 1992, Chap. 6 and 9). This procedure has proven robust for a variety of tasks, such as branch length estimation (Felsenstein, 1981), unbiased variance components estimation (Housworth et al., 2004; Ives et al., 2007), or ancestral state reconstruction (Royer-Carenzi and Didier, 2016). Under Gaussian models, the REML procedure is generally carried out using phylogenetic independent contrasts (Felsenstein, 1973).

By construction, for symmetrical processes such as the Brownian motion and the Cauchy process, marginalizing over the root value of a crown tree amounts to consider the likelihood of the process on the stem tree obtained using the following procedure: (i) first merge together the two branches starting at the root of the original tree, which gives us an unrooted tree, then (ii) pick one of its tip and root this unrooted tree at the parent of this tip, (iii) use the tip value as the starting value, and the tip branch as the stem branch, and finally (iv) discard the tip. Because of the symmetry assumption, any tip chosen in this procedure results in the same likelihood. It follows that REML procedures for the estimation of the dispersion parameter and the reconstruction of ancestral states and increments on crown trees under the CP can be straightforwardly derived from the maximum likelihood (ML) case of stem trees.

Note that, in theory, because of the symmetry assumption, any tip can be selected to carry on these computations, leading to the same results. In practice, to improve the stability of our computations and get trees that are more balanced, we select the tip that is “the closest” to all others on the tree, that is the tip *a* such that 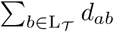 is minimal, with *d*_*ab*_ the distance on the tree between *a* and *b*.

### 2.4 Practical Implementation

We implemented the likelihood and density computation algorithms in the `C` language to improve speed, with an interface in `R` through the package `cauphy`. We used the `nloptr` optimization package (Johnson, 2021), with the Multi-Level Single-Linkage (Rinnooy Kan and Timmer, 1987; Kucherenko and Sytsko, 2005) and BOBYQA (Powell, 2009) algorithms for global and local numerical optimization. For the initialization of the parameters, we used robust estimates that ignored the phylogenetic correlations, but were relatively close to optimal parameters in practice. Namely, we normalized the trait values by dividing them by their associated tip heights, and then used the median for the root value and the *Q*_*n*_ statistics from Rousseeuw and Croux (1993) as an alternative to the inter-quartile range (IQR) or Median of Absolute Deviations (MAD) for the dispersion.

## 3 Simulation Studies

### 3.1 Model Selection

#### 3.1.1 Setting

To test our ability to distinguish gradual from pulsed evolution using the CP, we simulated the evolution of a trait on a tree using various models, and tried to recover the true underlying model from datasets made of the resulting tip trait values. In all simulations, we used the empirical tree from the Lizard dataset (Mahler et al., 2013, see below), normalized to unit height. In the following, a “dataset” hence consists of this fixed tree, and of a vector of trait values at its tips. We simulated a first dataset according to a simple BM, with root fixed to 0, and variance 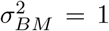. As the CP does not have a variance, we used the population median absolute deviation (MAD) to produce datasets with similar characteristics. It is well known that the MAD of a standard Gaussian is equal to its third quartile 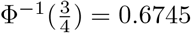, and that the MAD of a centered Cauchy variable is equal to its dispersion parameter (see e.g. Rousseeuw and Croux, 1993). We hence simulated a second dataset according to a CP with dispersion parameter *σ* = 0.6745, so that it had the same MAD as the standard Brownian dataset. To see if stabilizing selection or explosive change could lead to the spurious selection of a CP, we simulated two datasets using an Ornstein-Uhlenbeck (OU; Hansen, 1997) or and Early Burst (EB; Harmon et al., 2010) process. For the OU, we took the phylogenetic half-life of the process to be 20% of the tree-height, keeping a stationary variance of 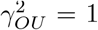, so that the selection strength was equal to *α*_*OU*_ = 3.47 and the process variance to 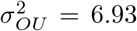. Exploiting the similarities between the OU and the EB (Uyeda et al., 2015; Aristide et al., 2018), we took and Early Burst with the same variance 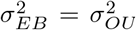, and opposite rate *r*_*EB*_ = *− α*_*OU*_. To assess how sensitive the CP is to various levels of pulsed evolution, we also simulated data according to a Normal Inverse Gaussian (NIG; Barndorff-Nielsen, 1997) process. The NIG process is also a pure jump Lévy process, and it has already been used in the literature to model trait evolution (Landis and Schraiber, 2017). In addition to a location and scale parameters *μ*_*NIG*_ and *δ*_*NIG*_, the symmetric NIG distribution has a tail parameter *α*_*NIG*_, larger values of *α*_*NIG*_ implying lighter tails (Hanssen and Oigard, 2001). Interestingly, when *α*_*NIG*_ = 0, the NIG reduces to a Cauchy distribution, with the same location and scale parameters, and, when *α*_*NIG*_*→ ∞*, with *δ/α*_*NIG*_ constant equal to *σ*^2^, the NIG converges to a Gaussian distribution with variance *σ*^2^ and same location (Barndorff-Nielsen, 1997). This *α*_*NIG*_ parameter can hence be used to produce datasets that have patterns of variation in between the BM and CP models. We simulated datasets according to the NIG process, all with the same MAD value equal to 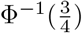 as before, and with an excess kurtosis varying from 1 to 10 (larger kurtosis being associated with lower *α*_*NIG*_ values). All the parameters used in this simulation study are summarized in Table 1.

**Table 1:**
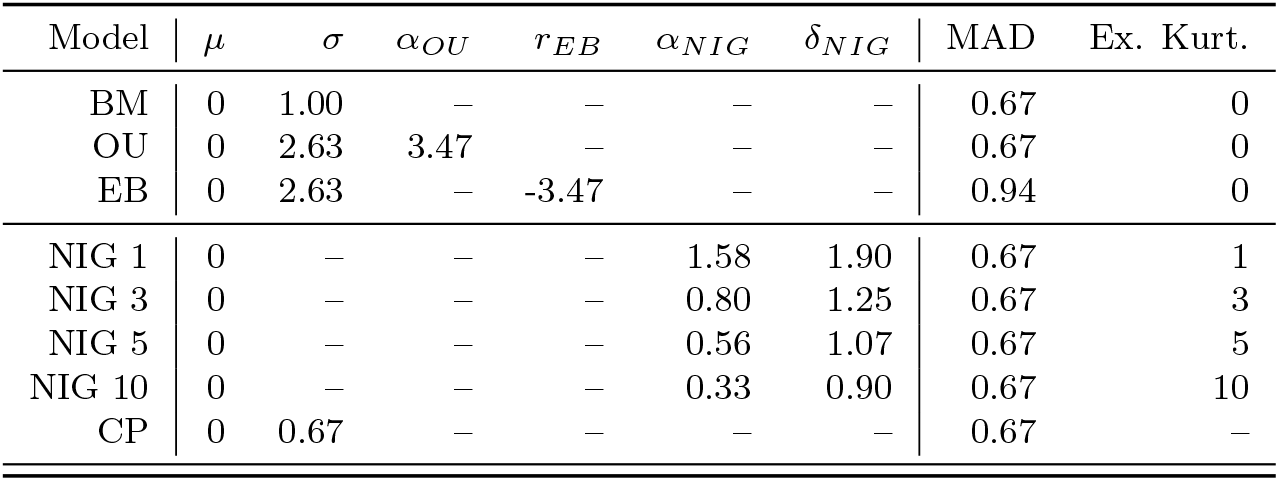
Parameters used for the simulation of the processes on the Lizard tree. *σ* is the standard error parameter for Gaussian processes, and the dispersion parameter for the CP. *μ* is the starting value. The population MAD is the same for all processes, except for the EB which parameters were taken from the OU. The excess kurtosis for Gaussian processes is 0, varies from 1 to 5 for the NIG processes, and is not defined for the CP.

| Model | $\mu$ | $\sigma$ | $\alpha_{OU}$ | $r_{EB}$ | $\alpha_{NIG}$ | $\delta_{NIG}$ | MAD | Ex. Kurt. |
| --- | --- | --- | --- | --- | --- | --- | --- | --- |
| BM | 0 | 1.00 | — | — | — | — | 0.67 | 0 |
| OU | 0 | 2.63 | 3.47 | — | — | — | 0.67 | 0 |
| EB | 0 | 2.63 | — | -3.47 | — | — | 0.94 | 0 |
| NIG 1 | 0 | — | — | — | 1.58 | 1.90 | 0.67 | 1 |
| NIG 3 | 0 | — | — | — | 0.80 | 1.25 | 0.67 | 3 |
| NIG 5 | 0 | — | — | — | 0.56 | 1.07 | 0.67 | 5 |
| NIG 10 | 0 | — | — | — | 0.33 | 0.90 | 0.67 | 10 |
| CP | 0 | 0.67 | — | — | — | — | 0.67 | — |

We simulated 100 datasets of tip trait values under each model. On each dataset, we used the function `phylolm` (Ho and Ané, 2014a) to fit the the Gaussian processes (BM, OU with fixed root, EB), both with or without error, and the function `fitCauchy` from our `cauphy` package to fit the CP. We used the maximum likelihood framework (with fixed root) for each fit, and selected the best model using the simple AIC score (Akaike, 1974).

We further replicated these analyses twice, first using a larger variance (from 1 to 100) for the reference BM while keeping homogeneous MAD values for all processes, and second adding an independent Gaussian noise at the tips of the tree to the trait resulting from each process, with an error variance going from 0.01 to 10 (see Supplementary Material Section C).

#### 3.1.2 Gaussian Processes are not Mistaken for Cauchy Processes

When the dataset was simulated under a Gaussian process (BM, OU or EB), the AIC always selected a Gaussian process, with the right model being chosen in the large majority of cases, confirming that there was a good recovery power also among Gaussian processes (see Fig 4). The other way around, when the dataset was simulated according to a CP, the AIC always selected a CP. Changing the AIC score to an AICc (Sugiura, 1978) or a BIC (Schwarz, 1978) did not qualitatively change the results in this simulation setting, that has a large number of samples (Fig S1). These results show that the signature of the CP is very different from the signature of a Gaussian process, and that there is a very high segregation power between these two modes of evolution.

**Figure 4:**
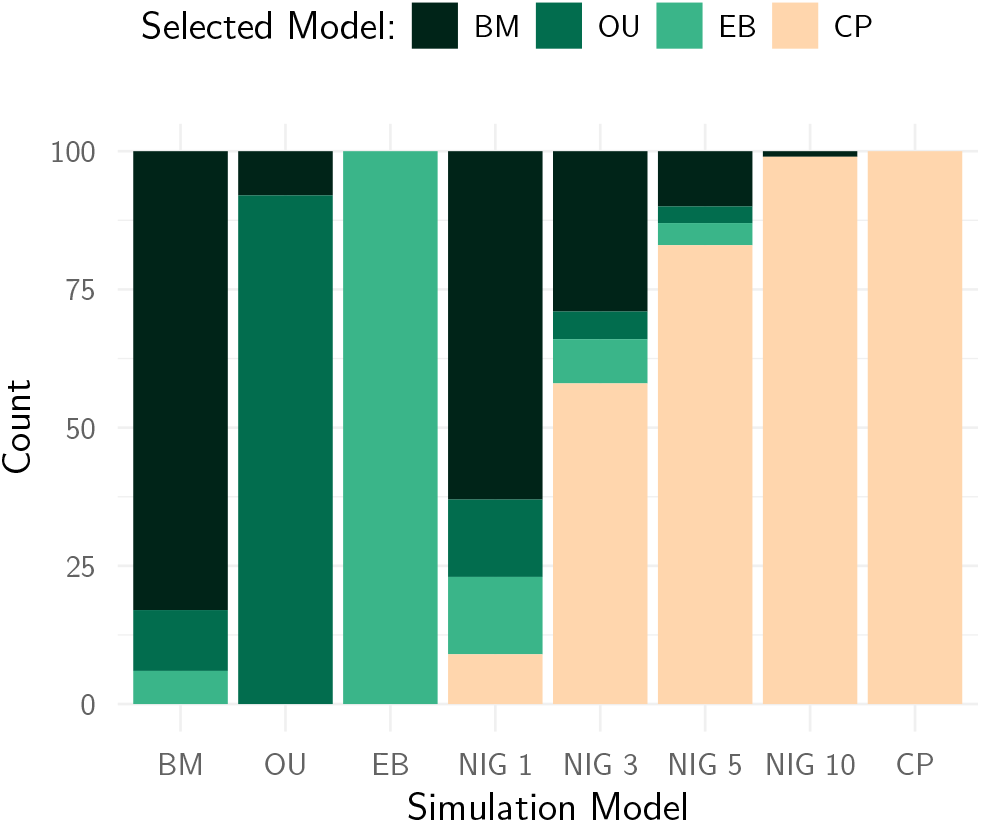
Model selection results using the AIC for 100 datasets simulated on the Lizard tree (Mahler et al., 2013), with various processes. Simulation processes are on the x axis, and include three Gaussian processes (BM, OU, EB), the NIG process with various levels of excess kurtosis (from 1 to 10), and the Cauchy process. All simulation processes but the EB have the same MAD (see Table 1). For each simulation model, the proportion of time each fitting model is selected is shown as a bar plot.

Considering models with higher variance or dispersion did not change these results (Fig. S6). However, additional independent Gaussian noise with high variance impaired model selection, and, as expected, favored the BM (with measurement error) over all other processes. Data simulated from the CP however remained partially identifiable, even in the presence of a high noise level (Fig. S7).

#### 3.1.3 The Cauchy Process Captures Pulsed Evolution

When the data was simulated according to a NIG process, the true simulation model was not included in the fitting models. From Figure 4, we can see that the Cauchy process is largely preferred over Gaussian processes for the NIG datasets, as soon as its excess kurtosis is large enough, that is, when the NIG goes away from the BM. As the NIG is a pure jump process that can also model non-gradual evolution (Landis and Schraiber, 2017), this result shows that the CP is able to pick up a different kind of pulsed evolution.

### 3.2 Parameter Inference

#### 3.2.1 Setting

We assessed the quality of the ML and REML estimators of the dispersion and root position parameters of a CP on a tree using simulations. We simulated pure birth trees conditioned on having from 10 to 250 tips using package ape (Stadler, 2011; Paradis, 2015), that we re-normalized to unit height. We then simulated trait data on each tree using a Cauchy process with dispersion 1 and root value 0. Each configuration was replicated 500 times. We then fitted a CP on each dataset, using either ML (with fixed root) or REML, and compared the estimated values of the parameters with the true simulation values. For the REML, we used ancestral trait reconstruction to get the posterior trait density at the root on a grid of values between *−*10 and 10 with step 0.01, and constructed a 95% highest posterior density interval (HDPI) using R package `HDInterval` (Meredith and Kruschke, 2022).

#### 3.2.2 Estimation of the Dispersion

Figure 5 shows that the method is able to correctly estimate the dispersion parameter for various tree sizes. When the tree is small (less than 50 tips), the ML method tends to under-estimate the dispersion, while the REML estimate seems to be un-biaised. This behavior matches similar findings in the Gaussian phylogenetic case (Housworth et al., 2004; Ives et al., 2007). For larger trees, the REML estimate also seems to have a slightly lower variance than the ML. Each fit took less than 10 second to compute on a standard laptop computer (MacBookPro 13-inch, M2, 2022), usually taking less than 1 second from trees with 100 tips or less (see Supplementary Figure S2). The REML fit, that only maximizes over one parameter instead of two, was consistently faster than the ML fit. When the tree had more that 200 tips, we observed that the parameter estimation tends to deteriorate. This phenomenon could be explained by the numerical instabilities of our algorithm (see Discussion), that can become unreliable for larger trees.

**Figure 5:**
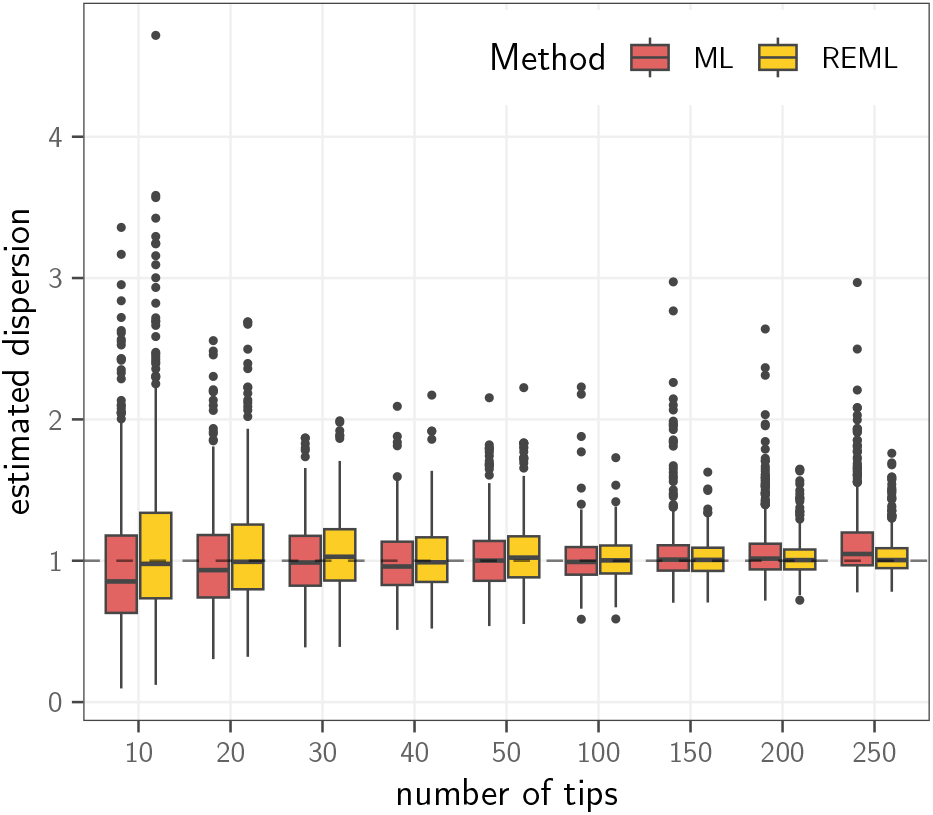
Estimation of the dispersion parameter of a CP on pure-birth trees of growing sizes (x axis), using the maximum likelihood with fixed root method (ML, dark red) or the restricted maximum likelihood (REML, light orange). The true simulation value is indicated as a dashed line. Each boxplot is on 500 replicates.

#### 3.2.3 Estimation of the Root Value

Figure 6 shows that the estimation of the root value was unbiaised, both in the ML or REML frameworks, for any size of tree. A small fraction of 5.5% of the datasets over all scenarios produced point estimates that had an absolute value larger than 2, far away from the true value of 0. These could be cases where, because of the jumping nature of the Cauchy process, the root value gets “forgotten” along the tree. However, Figure 7 shows that, in the REML case, the empirical coverage of the HDPI (fraction of simulation where the true value lies in the estimated interval) is close to its nominal rate of 95%, for all sizes of tree.

**Figure 6:**
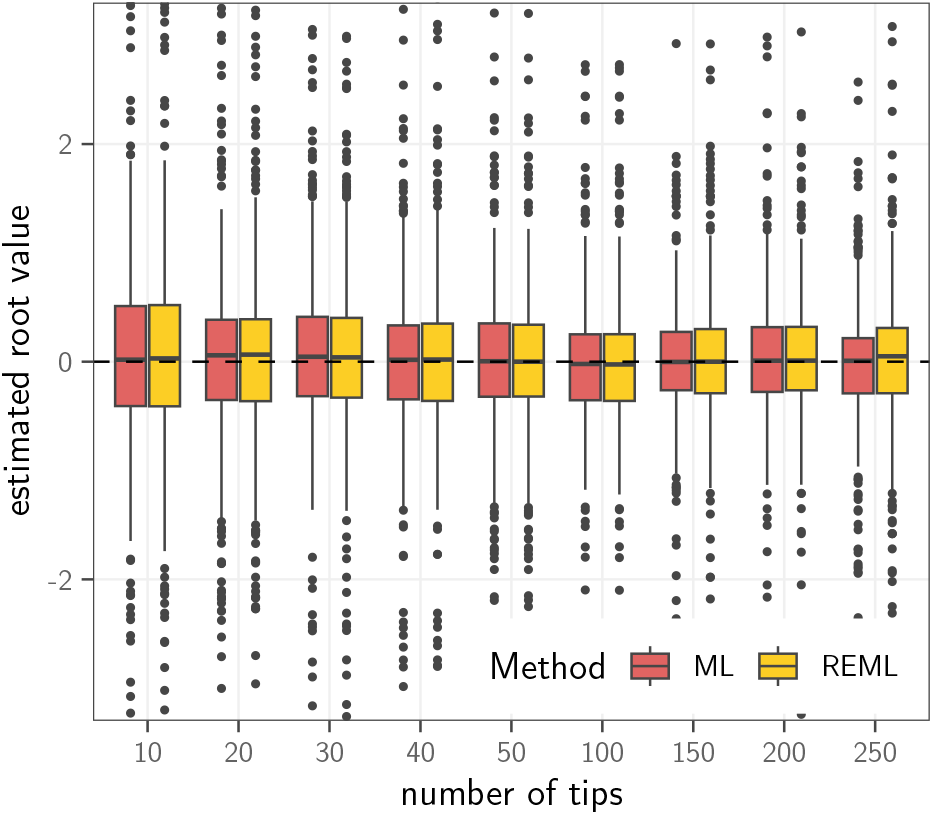
Estimation of the root trait value of a CP on pure-birth trees of growing sizes (x axis), using the maximum likelihood with fixed root method (ML, dark red) or the restricted maximum likelihood (REML, light orange). Each boxplot is on 500 replicates. Outlier estimates with absolute value larger that 3 (3.1% of the points) are not shown on the graphic.

**Figure 7:**
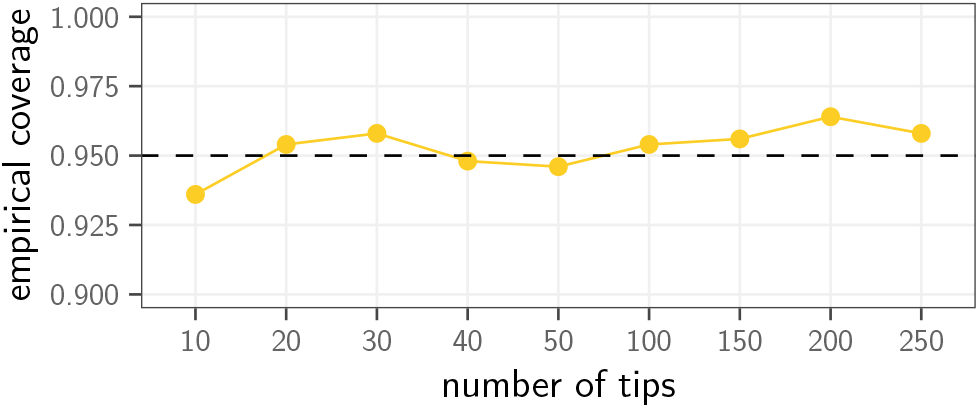
Fraction of simulations where the true simulation value for the root lies in the 95% highest posterior density interval computed from the ancestral posterior reconstruction of the root trait value using the REML (empirical coverage over 500 replicates for each tip number).

## 4 Empirical Examples

### 4.1 Greater Antilles Lizards

#### 4.1.1 Data

The adaptive radiation of anoles in the Greater Antilles is well documented (Losos, 2009), and has been hypothesized to follow a Simpsonian “evolution by jumps” (Mahler et al., 2013). Previous analyses of this dataset includes models based on Brownian Motion or Ornstein-Uhlenbeck with shifts (Mahler et al., 2013; Eastman et al., 2013; Bastide et al., 2018), or Lévy processes (Duchen et al., 2017). We considered here the dataset from Mahler et al. (2013), that consists of 100 lizard species on a time-calibrated maximum clade credibility tree, with each species associated with an ecomorph. As in Duchen et al. (2017), we focused our analysis on the log-transformed snout-to-vent length (SVL), which is highly correlated with all the other morphological measurements (Bastide et al., 2018).

#### 4.1.2 Methods

We fitted a Cauchy process on this dataset, using the ML or REML method. Using the AIC criterion (Akaike, 1974), we compared these results with fits obtained using traditional Gaussian processes, such as the Brownian Motion (Felsenstein, 1973), the Ornstein-Uhlenbeck (Hansen and Martins, 1996) and the Early Burst (Blomberg et al., 2003; Harmon et al., 2010), all fitted with `phylolm` (Ho and Ané, 2014a) (ML) or `Rphylopars` (Goolsby et al., 2017) (REML). We then carried an ancestral state reconstruction using the Cauchy REML fit, on a grid of 100 values between 3 and 5.5 for all trait values at internal nodes, and between *−*1.5 and 1.5 for all increments on branches.

#### 4.1.3 Model fit

Compared to Gaussian processes, the Cauchy process increased the log-likelihood by about 8 units, and had a much smaller AIC (Table 2). Consistent with a pulsed evolution hypothesis, the CP was hence clearly favored over any Gaussian process.

**Table 2:** Fits of Gaussian (BM, OU and EB) and CP models on the lizard dataset from Mahler et al. (2013), with 100 species, and using the snout-to-vent length (SVL) trait measurement. The first two lines are the (restricted) log likelihood and AIC scores for the ML (left) and REML (right) fits. The parameters are *μ* for the ancestral root value (ML), *σ* for the Gaussian standard error or Cauchy dispersion, and *θ* represents the selection strength *α* for the OU, and the rate *r* for the EB. The computation time for each fit on a standard laptop is given in seconds.

|  | ML |  |  |  | REML |  |  |  |
| --- | --- | --- | --- | --- | --- | --- | --- | --- |
|  | BM | OU | EB | CP | BM | OU | EB | CP |
| logLik | -4.70 | -4.70 | -4.29 | **4.44** | -6.00 | -6.01 | -5.36 | **3.10** |
| AIC | 13.40 | 15.40 | 14.57 | **-4.88** | 14.01 | 16.01 | 14.73 | **-4.21** |
| $\mu$ | 4.07 | 4.07 | 4.07 | **4.00** | — | — | — | — |
| $\sigma$ ($\times 10^{-3}$) | 52.18 | 52.18 | 68.34 | **4.51** | 52.45 | 52.45 | 73.48 | **4.48** |
| $\theta$ | — | 0.00 | -0.01 | — | — | 0.00 | -0.02 | — |
| time (s) | 0.02 | 0.11 | 0.01 | **1.55** | 0.01 | 0.25 | 0.05 | **1.36** |

#### 4.1.4 Ancestral Reconstruction

As in Duchen et al. (2017), we found evidence for large positive increments on branches leading to clades with species from the “crown giant” ecomorph, that have particularly large body sizes (Fig. 8). The clade highlighted with bold edges on Figure 8 is particularly interesting, as it exhibits bi-modal posterior trait and increment distributions, that can support two possible hypotheses about body size evolution of these species. One possible scenario is consistent with the one exhibited in Duchen et al. (2017), with medium sized ancestors (“orange” mode on ancestral nodes), and shifts towards large values on branches leading to species from the “crown giant” ecomorph, and to the clade of “false chameleons”, namely *A. guamuhaya, A. porcus, A. chamaeleonides* and *A. barbatus*, from the former genus *Chameleolis*, that are known to have unique characteristics. However, an other scenario that is supported by this dataset would see a shift toward large body sizes at the crown of the clade leading to large ancestors (“purple” mode on ancestral nodes), and shifts towards smaller values on branches leading to the two smaller species of *A. christophei* and *A. eugenegrahami*.

**Figure 8:**
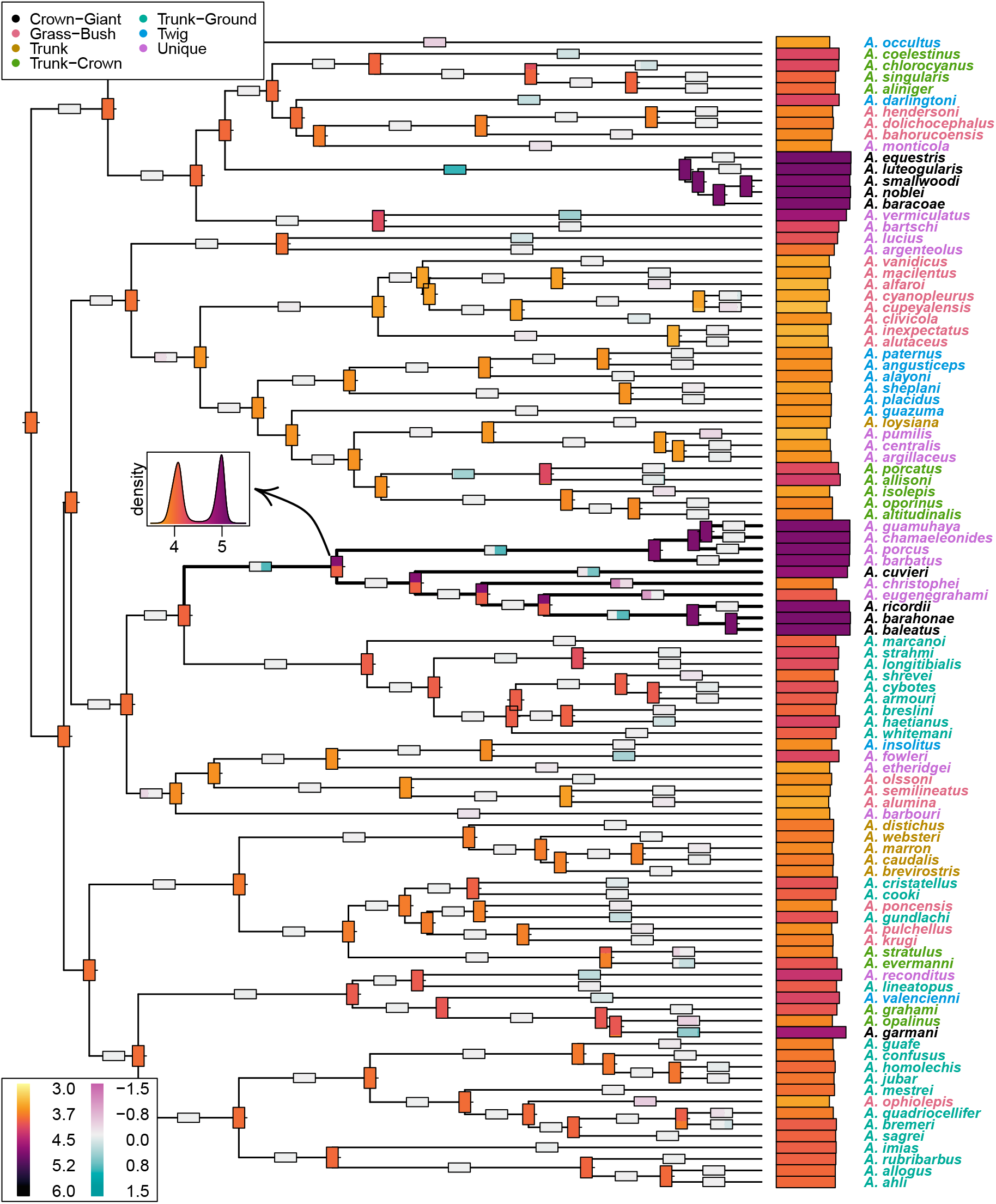
Ancestral trait reconstruction for the lizard dataset from Mahler et al. (2013), with the log-transformed snout-to-vent length (SVL) trait measurement, using the Cauchy REML model fit. Species names are colored according to their ecomorphs. Measured traits are shown as a bar plot at the tips of the tree, with a color gradient encoding the trait value, from light yellow (small) to dark purple (large). Modes of the posterior trait reconstructions are showed at internal nodes, with a width proportional to the relative height of the modes (nodes with only one color have a single mode), and the same color scale as tip values. Modes of the posterior branches increments are showed on the edges, using the same convention, but with a diverging color scale so that increments close to zero are white, positive increments are blue, and negative increments are red. The clade discussed in the text is highlighted in bold. The posterior density for the ancestral reconstruction of the stem of this clade, which is bi-modal, is shown as an inset plot, colored with the same gradient.

#### 4.1.5 Timing

The initial Cauchy fit was fast, taking less than 2 seconds for the CP (including the root state estimation in the ML case). The additional character reconstruction took around 2 seconds total for all the 99 ancestral nodes (using a single thread), and around 17 seconds total for all the 198 edges (using six threads in parallel). All analyses were carried on a standard laptop computer (MacBookPro 13-inch, M2, 2022).

### 4.2 West Nile Virus

#### 4.2.1 Data

The epidemic of WNV in America, from its first detection in New York city in 1999 to its spread in California and Mexico in the early 2000s, caused more than 1,200 deaths in the United States and was closely monitored by the health authorities. Pybus et al. (2012) studied the geographical spread of the virus in an integrated framework using BEAST (Suchard et al., 2018), with a dataset of 104 whole genomes collected between 1999 and 2007 along with the sampling coordinates (latitude and longitude). We reproduced the analysis using the best fitting model they selected, which consisted for the nucleotides of a general time-reversible (GTR) substitution model with a gamma distribution of among-site rate heterogeneity, an uncorrelated lognormal relaxed molecular clock and a Bayesian skyline coalescent model prior, and for the continuous locations of a relaxed random walk (RRW) with a gamma prior for the mixing rates, and an added uniform jitter with window size 0.5 applied to duplicated locations. We followed the instruction from the associated BEAST workshop tutorial (available at https://beast.community/workshop_continuous_diffusion_wnv) to set up the analysis, and ran the MCMC chain for 250 million states, sampled every 50,000 states, using BEAST v1.10.5 (prerelease #dc4f12c). The convergence of the chain was assessed with `Tracer` (Rambaut et al., 2018), and we obtained a maximum clade credibility (MCC) tree using `TreeAnnotator`, with times summarized with the common ancestor heights option to avoid negative branch lengths. In all subsequent analyses, we used the MCC tree and the jittered continuous locations as the primary dataset.

#### 4.2.2 Methods

We fitted a CP using our framework on both the latitude and longitude independently, and reconstructed ancestral nodes an increments on all branches using a grid of 1000 values. We then compared the fit of the CP with other Gaussian models fitted with phylolm using the AIC score. Using the Hamiltonian Monte Carlo (HMC) framework developed in Fisher et al. (2021), we also fitted a RRW of the two traits, with correlations, and an inverse gamma prior with parameter 1*/*2, referred to as a “Cauchy RRW” in the literature, but that is distinct from a true CP (see above). To compare the two processes, we also fitted two independent “Cauchy RRW” on the latitude and longitude using BEAST, and a CP on the same continuous data, but using a tree with square rooted branch lengths. For maximum likelihood fits, confidence intervals (CI) were obtained using the approximate Hessian, and for Bayesian fits, we used the highest posterior density interval (HDPI) as implemented in the R package `HDInterval` (Meredith and Kruschke, 2022). All chains were run for 500,000 states, with parameters and ancestral nodes traits sampled every 500 states. We used a vague exponential prior with mean 1 and an uniform prior between

*−*1 and 1 for the variance and correlation parameters of the RRWs.

#### 4.2.3 Model fit

Compared to Gaussian models, the CP has a much higher log-likelihood, and is selected with an AIC at least 60 units better for both the latitude and longitude compared to the EB model, which was the second best model (see Supplementary Table S1).

#### 4.2.4 Comparison with RRW

The independent “Cauchy RRW” and CP on the square-root re-scaled tree gave very close estimated of the dispersion parameter of the latitude and longitude traits, for both the point estimates and confidence intervals (see Table 3). Up to numerical differences related to the fitting method (maximum likelihood versus Bayesian MCMC with vague priors), this result is consistent with the theoretical equivalency of these models.

**Table 3:**
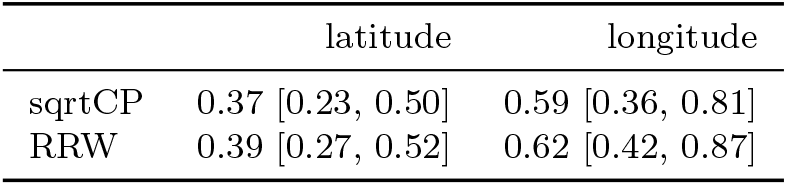
Comparison of dispersion estimates with the Cauchy process on the tree with square-rooted branches (sqrtCP) and the relaxed random walk with a “Cauchy” rate on the priors (RRW), both fitted independently on the latitude and longitude data of the WNV MCC tree.

#### 4.2.5 Long-distance Dispersion Events

As in Pybus et al. (2012), we recovered several rapid and long-distance dispersion events, that is allowed by the jumping nature of the CP (see Supplementary Figure S4). In particular, the first north to south travel of the virus allows us to compare the RRW and CP approaches (see Fig. 9). Indeed both models exhibit the same general pattern, with a rapid shift from New-York to Florida in the first half of 2000, on the small branch going from the black to the dark orange node. However, the CP also supports an alternative hypothesis, where the shift does not occur on this branch, but latter on the tree after the light yellow node, corresponding to a scenario where the virus first spreads to Ohio, and then to Florida, instead of radiating to back to Ohio from Florida as in the first scenario. This alternative scenario involves shifts of lower size, and its absence in the RRW case could be related to the fact that the RRW is a CP on a square-rooted tree, and hence has a larger dispersion parameter than the standard CP for short branches of length less than one, possibly allowing for longer dispersion events. As the RRW density is based on an MCMC, it could also be possible that one of the modes was never sampled in the chain. Finally, the RRW takes some correlations between the traits into account, while our CP treat them as independent, which could also be a source of discrepancy.

**Figure 9:**
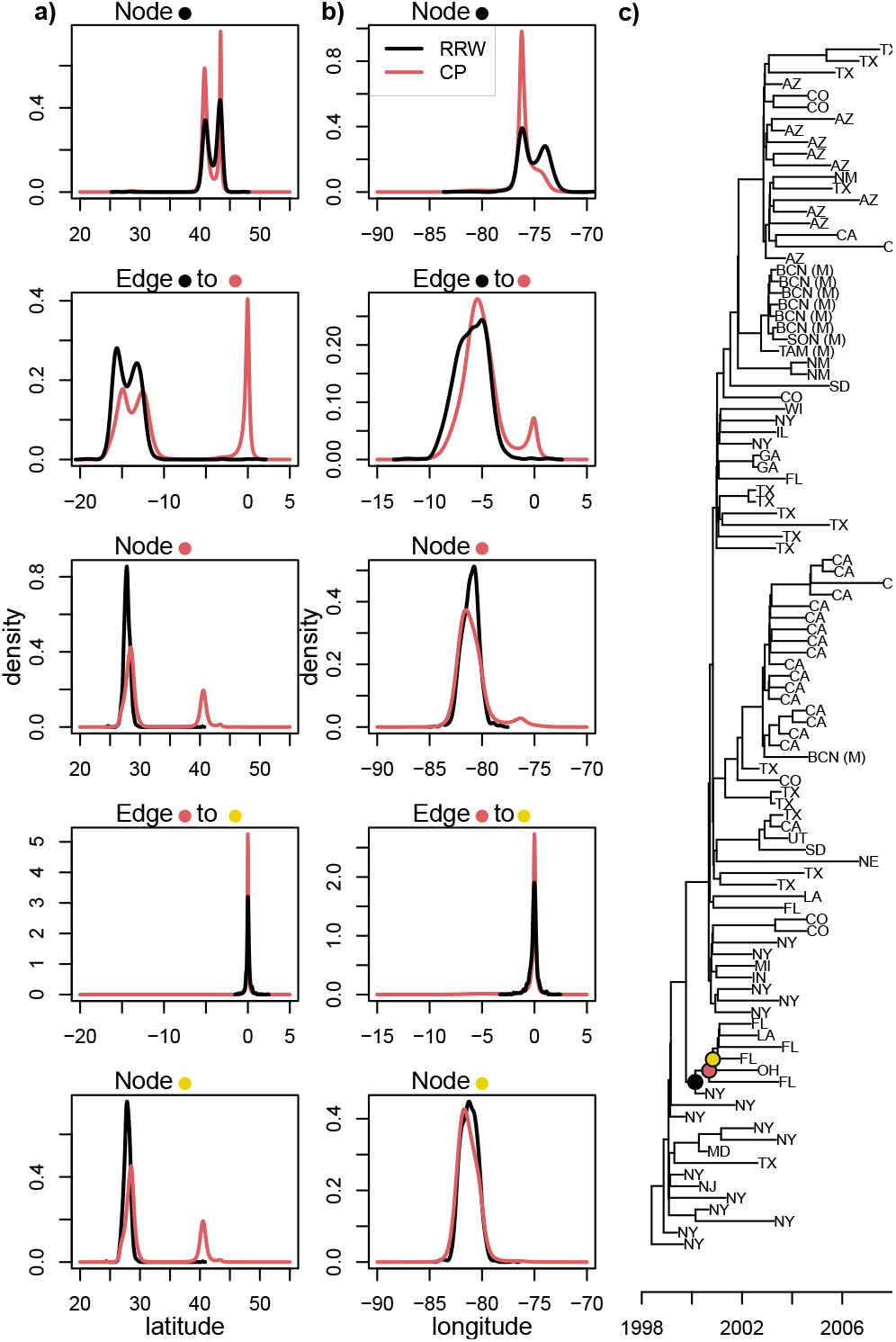
Ancestral node and increment reconstructions for the latitudes (**a**) and longitudes (**b**) using the “Cauchy RRW” (black curves) and CP (dark orange curves) on the MCC tree of the WNV (**c**), for the nodes and edges marked with colored dots on the tree. For the RRW, curves are kernel density estimates from states sampled in the MCMC chain.

## 5 Discussion

### 5.1 The Cauchy Process for Trait Evolution

#### 5.1.1 The Cauchy Distribution

Various situations may lead to observe the Cauchy distribution in practice. For instance, it arises as the ratio of two independent normal Gaussian random variables, so that it could be well suited for some biological measurements, such as in the study of morphological evolution, where scores can be defined as ratios of several osteological measurements (Schnitzler et al., 2017). Further, each stable distribution enjoys an analogue of the central limit theorem, and is the limiting point of a whole domain of attraction in the space of distributions (see e.g., Klenke, 2014). In the Gaussian case, this makes the Brownian Motion a natural model for many traits, such as polygenic characters under neutral evolution, or adaptive traits in a randomly changing environment (Felsenstein, 2004). The Cauchy domain of attraction, although fully characterized in theory, is not as simple as the Brownian one. An interesting special case is given by Shapiro (1975). Assume that *X*_1_, …, *X*_*n*_ is an independent identically distributed sequence of random variables, with common probability density function *f*, such that *f* is continuous in 0 with *f* (0) *≠* 0 and *f* derivable with *f*^*′*^ bounded in the neighborhood of 0. Then the empirical mean of the *inverses*, 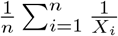, converges in distribution towards a Cauchy random variable, with a given position parameter, and dispersion *σ* = *πf* (0). This simple analogue of the central limit theorem for inverse effects, as well as a deeper study of its general domain of attraction, could help us to gain some future insights on the biological phenomena that could be effectively modeled as Cauchy processes.

#### 5.1.2 The Pure Jump Cauchy Process

As a pure jump process, we showed that the CP could capture pulsed evolution phenomena. Simulation studies demonstrated that there was a strong segregation signal between Gaussian and Cauchy processes, that were never mistaken from one another, and that the CP could detect other kinds of jumping patterns, such as the NIG. The empirical Lizards dataset study illustrated that, using a model that has as many parameters as the standard BM, and that is very efficient to fit, we could recover some of the main conclusions of Duchen et al. (2017).

#### 5.1.3 The Relaxed Random Walk Cauchy Process

The CP can be re-framed as a scaled relaxed random walk, where each branch has its own variance parameter, drawn from a gamma distribution with parameters scaled by the branch length. We emphasize here that this process is different from the classical RRW of Lemey et al. (2010), and implemented in BEAST (Suchard et al., 2018), where the branch variance variables are identically distributed. This latter model cannot be associated to any trait evolution process on the tree, as opposed to the CP, which has a nice jump process interpretation. We showed that it could be described as a CP, but on a time transformed tree, where the square root of each branch length is taken. For trees with branches smaller than one, this leads to longer branches in the transformed tree, and hence possibly more jumping events than expected under a simple CP, as we observed for the WNV dataset. However, this empirical example showed that, in a maximum likelihood framework, we could use the CP to study the geographical spread of a virus, and recover the main conclusions from previous Bayesian RRW studies. These results are encouraging, and show that the CP can be a cheap and versatile alternative to complex jumping or parameter rich models currently in use in the literature.

### 5.2 Ancestral Reconstruction and Shift Detection

#### 5.2.1 Ancestral Reconstruction

Using the CP to model trait evolution, we derived recursive algorithms to explicitly compute the posterior distribution of trait values at internal nodes, conditionally on tip values. Looking at the root state, we showed that this reconstruction was accurate, with the empirical coverage of HDPI close to their nominal values for data simulated according to a CP on birth-death trees of various sizes. It would be interesting to compare our Cauchy reconstructions with other reconstruction techniques, either Gaussian (Royer-Carenzi and Didier, 2016), or non-Gaussian (Elliot and Mooers, 2014; Gao and Wu, 2022b). We expect our reconstructions to be more accurate than Gaussian ones for jumping processes, while being more numerically efficient than currently existing methods that can accommodate for more complex Lé vy processes.

#### 5.2.2 Identifiability of Evolutionary Scenarios

On a toy example as well as on both empirical examples, we illustrated how ancestral trait reconstructions could be multi-modal, and how this feature could be interpreted in terms of shifts in the trait value on the tree. Compared to shift detecting methods, that use Gaussian processes with fixed (but unknown) shifts on some branches (Ingram and Mahler, 2013; Khabbazian et al., 2016; Bastide et al., 2018), that are prone to identifiability issues (Bastide et al., 2017), the CP based reconstruction provides a nuanced picture, with the different modes associated to several possible evolutionary scenarios of the trait value. As the CP implicitly defines a distribution on the occurrence and magnitude of jumps, it can also provide us with an idea of which scenario is more probable given the data and the model, through the comparison of the weights associated with each modes.

#### 5.2.3 Shift Detection

The posterior distribution of the trait increment on the branches of the tree can confirm and extend the insights gained from the node reconstructions. We exhibited several instances where this increment was centered on a value different from 0, or had several modes, around 0 and other values. It is tempting to interpret these as signal for a shift, i.e. as a sign that the trait evolved particularly fast on a given branch. In this study, we limited ourselves to the description of the posterior distribution, leaving the interpretation to the user. It would however be interesting to see if these could be used as a basis for automatic shift detection. This could be done, e.g., using a statistical test to detect branches where the prior (un-conditioned) increment distribution is significantly different from the posterior (conditioned to the tip data) one. Such a method could help focus the attention of researchers to branches that seem particularly interesting, and for which a detailed reconstruction is needed.

### 5.3 Likelihood Computation and Numerical Robustness

#### 5.3.1 Likelihood Computation

One of the main contributions of this paper is an explicit recursive algorithm to compute the likelihood of the CP on a tree, as detailed in Proposition 1. This algorithm is the corner stone of our approach, and is used for parameter estimation, model selection and ancestral trait reconstruction alike. It involves the computation of many sums, and has a time complexity that is quadratic in the number of tips, as opposed to linear times needed in the Gaussian case (Ho and Ané, 2014a; Mitov et al., 2020). For all the trees of reasonable size that we tested in this study, the computation times remain relatively short, a maximum likelihood fit never taking more than ten seconds on a standard laptop computer (see Supplementary Figure S2). This make our method practical to use in many situations.

#### 5.3.2 Theoretical Limitations

Unfortunately, this algorithm has several limitations, both theoretical and practical. Looking at Equation (4), we can see that the second term of the sum might imply a division by zero for tips *a* and *b* such that *t*_*k*:*a*_ = *t*_*ℓ*:*b*_ and *y*_*a*_ = *y*_*b*_, i.e. for two tips that are simultaneously (i) at the same temporal distance from their least common ancestor, and (ii) with a same trait value. Although condition (i) is common, and in particular verified for ultrametric trees, condition (ii) is theoretically impossible, as the probability of observing two tips with the exact same continuous trait value is zero. However, on empirical datasets, where measures are subject to rounding approximations, it is not uncommon to encounter tips with equal trait values. For instance, the well known *Chelonia* dataset (Jaffe et al., 2011) has about 46% of its entries repeated, as body size are often reported up to the centimeter only. In such cases, our method cannot be applied directly. A simple solution could then be to add a reasonable amount of random noise to the tip measurements, to make sure that they are all distinct. Such a jitter is for instance often applied to continuous phylogeographic data in virology to account for sampling location uncertainty (Dellicour et al., 2022). A good knowledge of the biological system under study can help the practitioner choose this noise, that can also empirically compensate for intra-species variations or bias in the measurement process. In such cases, as our algorithm is relatively fast, several independent replicates of the noisy dataset can easily be generated in order to check the robustness of the analysis with respect to the added noise.

#### 5.3.3 Numerical Robustness

Besides this theoretical limitation, we observed some numerical robustness issues of our algorithms for large trees with more than 200 tips, which impacted parameter inference (see Fig. 5). This issue is caused by the many sums that need to be computed in our algorithms. These sums involve terms of arbitrary size, positive or negative, that may almost cancel out, and lead to precision errors. Despite our best efforts to use robust computational tools, because of the very structure of the summations, we could not further improve our implementation, even when using arbitrary precision summation techniques. We stress here that it is important to check, on biological datasets, that the numerical likelihood computations and optimizations behaved properly. As a first step, we propose in our package cauphy to plot the profile likelihood around the estimated parameters, that should be smooth and with a clear maximum (which is the case for both empirical examples presented here, see supplementary Figures S3 and S5). A new, more efficient decomposition of the likelihood on the tree might be needed to further improve the accuracy of its computation on larger trees.

#### 5.3.4 Composite Likelihood

An other, complementary approach would be to use an approximation of the likelihood, for instance assuming independence between contrasts, to get a composite likelihood (Varin and Vidoni, 2005). Landis and Schraiber (2017) developed such a clever composite (restricted) likelihood approach, that relies on canceling out covariances between well-chosen contrasts on the tree, and then making the approximation that these contrasts are independent to ease computations. Note that, as the Cauchy distribution does not have any moment, this technique could not be directly applied to the CP, but it could be adapted to this case, and would benefit from closed form formulas for the characteristic function integration at nodes, instead of having to resort to numerical integration as in Landis and Schraiber (2017). Composite likelihoods are usually easier and more efficient to compute, and yield unbiaised estimates of the parameters of the model. However, they cannot be used directly for model selection or confidence interval estimation, and some corrective terms, that are sometimes difficult to estimate, are needed for both these tasks (Varin et al., 2011). Studying the robustness and efficiency of composite likelihood approaches to study pulsed trait evolution could be the focus of future work.

### 5.4 The Cauchy Process for Robust Regression

#### 5.4.1 Robust Phylogenetic Regression

Phylogenetic regression uses a fixed root model, with the root value giving the fixed effects, and the trait evolution model giving the structure of the residual errors. In a Gaussian setting (see e.g., Ho and Ané, 2014a), this model can be written as **Y** = **X*β*** + ***ϵ***, with **Y** the vector of trait values at the tips of the tree, **X** a matrix of predictors, ***β*** a vector of coefficients, and ***ϵ*** a Gaussian vector of errors, centered and with variance *σ*^2^**V**^tree^, with **V**^tree^ the covariance matrix given by the trait evolution model (for the BM, 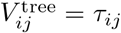 is the time of shared evolution between tips *i* and *j*). A similar regression model could be written in the Cauchy case, replacing the normal distribution of the errors by a multivariate Cauchy one, with position parameter 0, and structure given by a CP on the tree. Our algorithm can then be used to compute the likelihood of such a model, and we can resort to numerical optimization to infer the parameters ***β*** and *σ*. Such a model would amount to a Phylogenetic Cauchy Regression, which is known to be robust in the classical – “non-phylogenetic” – case (Liu and Tao, 2014). Methods for robust phylogenetic regression were recently proposed by Adams et al. (2022), with algorithms relying on the definition of *ad hoc* scores based on the S or MM procedures to replace the least squares. Our Cauchy framework could hence be seen as an alternative tool to perform robust phylogenetic regression, but with an explicit evolutionary model and using maximum likelihood estimation.

#### 5.4.2 Pagel’s lambda

The Pagel’s lambda (Pagel, 1999) transformation is frequently used in phylogenetic regression frameworks to relax the BM assumption. It relies on fitting a BM with variance 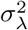, running on a modified tree with internal branch lengths multiplied by a parameter *λ*, and external branches lengthened so that tip heights remain unchanged. This heritability parameter (Housworth et al., 2004) can be used to test for phylogenetic signal (Revell, 2010), and, on an ultrametric tree with height *h*, this model is equivalent to a BM on the original tree with variance 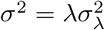 and added independent Gaussian noise at the tip with variance 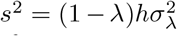 (Leventhal and Bonhoeffer, 2016). For any stable process, it is easy to see that this transformation is still valid, and have similar interpretations. In particular, a CP with dispersion *σ*_*λ*_ on a ultrametric tree with *λ* transformed branch lengths is equivalent to a CP on the original tree with dispersion *σ* = *λσ*_*λ*_ and added independent *Cauchy* noise at the tip with dispersion *s* = (1*−λ*)*hσ*_*λ*_. This transformation is straightforward to apply on any tree, and hence provides us with an easy way to accommodate for added independent noise in the data, and to relax the strict CP assumption.

### 5.5 Conclusion and Perspectives

In this work, we introduced the Cauchy Process on a tree, and showed that it was a versatile model for trait evolution, that could capture pulsed evolution phenomena, and provided the practitioner with an interesting and easy to fit alternative to the Brownian Motion and other Gaussian models. In addition to all the points discussed above, future work could involve refinements over the Cauchy process, for instance using a Ornstein-Uhlenbeck-Cauchy model (Garbaczewski and Olkiewicz, 2000) for noisy stabilizing selection, or extensions to multivariate traits.

## 6 Acknowledgements

We thank Stéphane Guindon, Denis Fargette, Pauline Rocu and Julien Clavel for insightful discussions; as well as Isabel Sanmartín, Michael Landis, Joshua Schraiber and two anonymous reviewers for their thoughtful comments. We have no conflict of interest to declare.

## 7 Supplementary Material

Supplemental information can be downloaded at https://doi.org/10.5281/zenodo.7802104. Data, including all the scripts used in simulations and empirical examples analyses, is available from the Dryad Digital Repository https://doi.org/doi:10.5061/dryad.d2547d86h, and the companion GitHub repository: https://github.com/pbastide/cauchy_paper. The `R` package `cauphy` is available at https://github.com/gilles-didier/cauphy/.

## Supplementary Material

### A Likelihood Computation

#### A.1 Cauchy Distribution

We consider Cauchy distributions with position parameter *μ* and dispersion parameter *σ ≥* 0. The density *f*_*C*_ and the associated characteristic function *ϕ*_*C*_ are

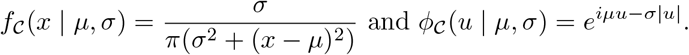

By applying the inverse Fourier transform to *ϕ*_*𝒞*_, the Cauchy density can be alternatively written as

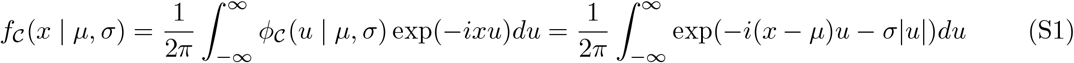

#### A.2 Joint Likelihood

We prove the following Proposition, which is a technical version of Proposition 1.

##### Proposition S1

(Likelihood Propagation). *Let 𝒯 be a stem tree with root r. The joint probability density of a vector of tip values* **y** *under the Cauchy model with dispersion σ and starting value μ can be written as:*

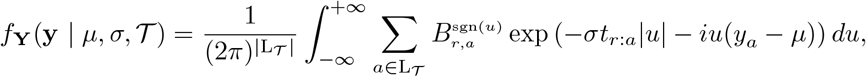

*where for all nodes n of 𝒯 and all tips a ∈ 𝒯*_*n*_, *t*_*n*:*a*_ *is the sum of the lengths of the branches of the path from n to a (length of the branch ending with n included) and the coefficients* 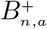 *and* 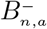 *are complex numbers recursively computed in the following way:*

*If n is a tip then* 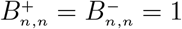

*If n is an internal node, with direct descendants k and ℓ:*

**–** *for all tips b ∈ 𝒯*_*k*_,

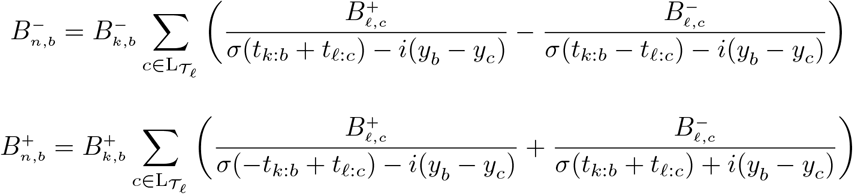

**–** *for all tips b ∈ 𝒯*_*k*_,

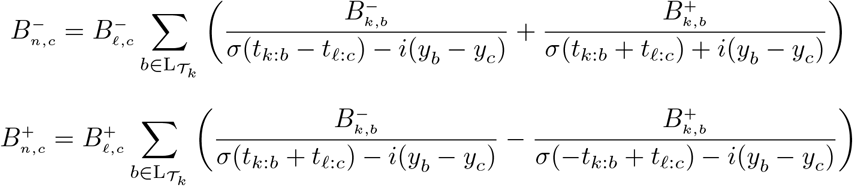

*Proof*. We prove this result by induction on the tree *𝒯* by considering partial likelihoods in a post-order, from the tips to the root. For any node *n* of *𝒯* and real value *μ*, we show that the partial likelihood of the vector of trait values 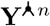 at the tips of sub-tree *𝒯*_*n*_ conditionally on the ancestor of *n* having trait *μ* is given by:

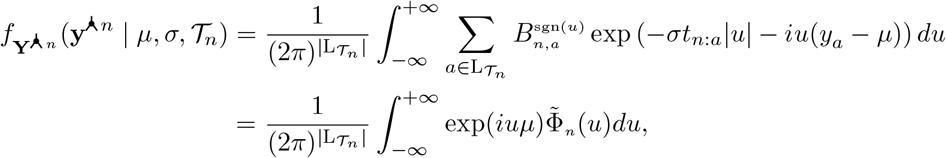

where we defined (dropping the dependency in *σ* and *𝒯* for simplicity),

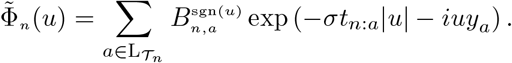

Note that *𝒯*_*n*_ is a stem tree, that includes the branch going from the ancestor of *n* to *n*. When *n* = *r*, the partial likelihood matches the full likelihood of the tree conditionally on the process starting at *μ* at the beginning of the branch ending at *r*.

If *n* is a tip, the partial likelihood 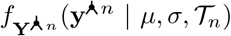 is the Cauchy density *f*_*𝒞*_(*y*_*n*_ | *μ, σt*_*n*_) of the branch ending at *n* with value *y*_*n*_, and Equation S1 gives us the result with 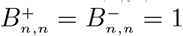.

Let now *n* be an internal node with *k* and *ℓ* its two direct descendants, and assume that the result holds for all *k* and *ℓ*. Using the propagation equation of the model and the conditional independence given by the tree, the joint probability density of the tips value vector 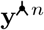 of the subtree *𝒯*_*n*_ is

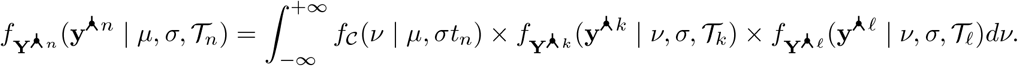

Using Equation S1 and by induction, we get:

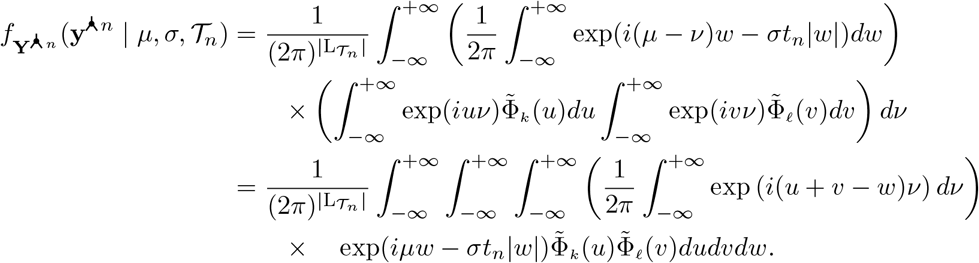

From Fourier analysis, we have that

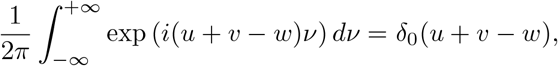

where *δ*_0_ is the Dirac mass at the origin (see e.g., Kammler, 2007, appendix). It follows that:

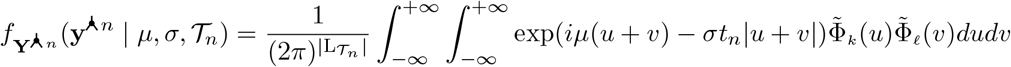

Let us consider the change of variables which replaces *v* with *w* = *u* + *v* and lets *u* unchanged. The determinant of the Jacobian associated to the transformation is 1, and the integral above becomes

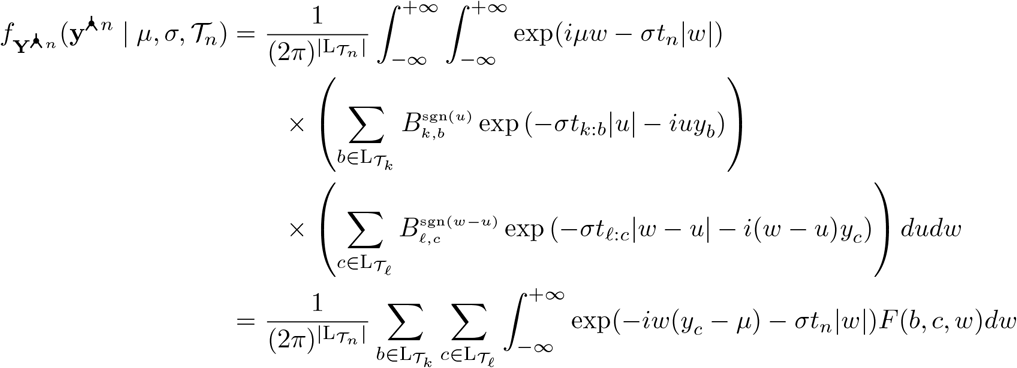

with

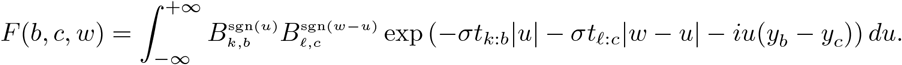

Integrating *u* over, we get, if *w ≥* 0,

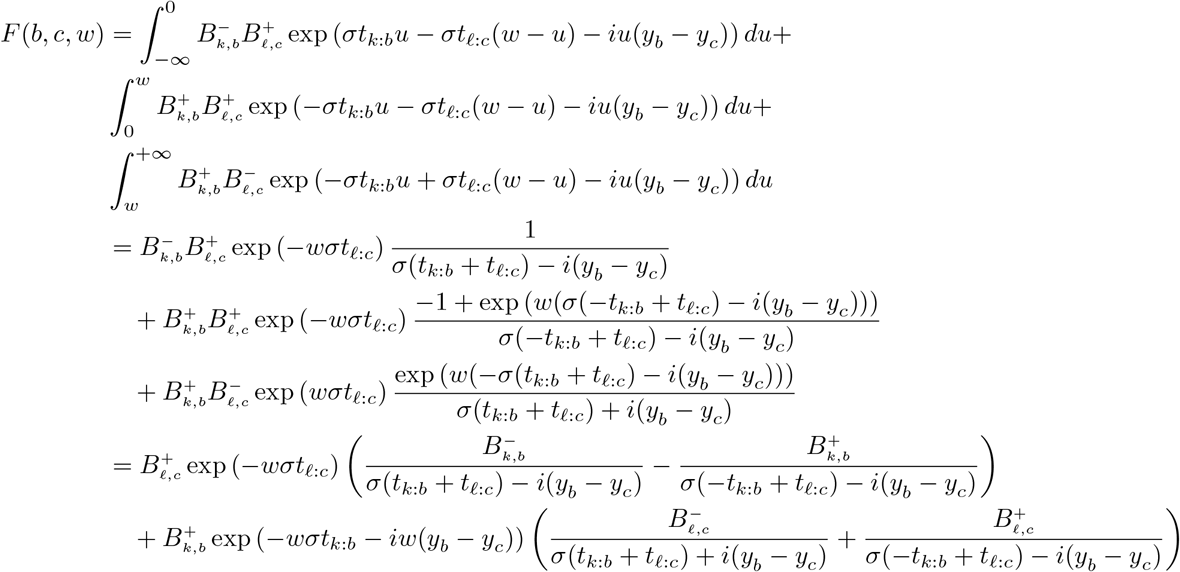

so that, using the recursion equations,

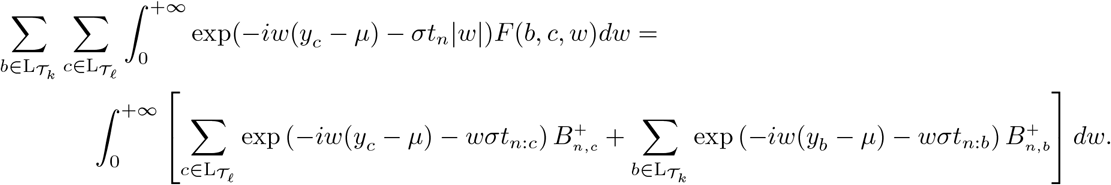

Symmetrically, for *w ≤* 0, we get

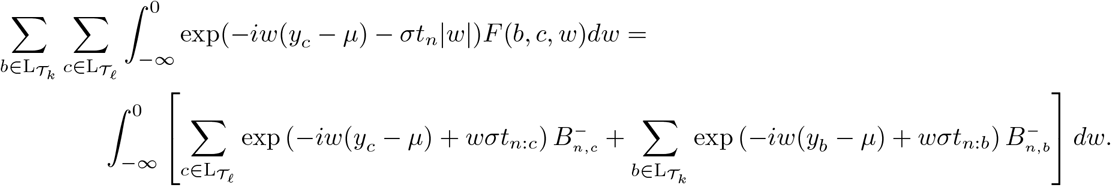

Joining both equations together, we get the expected expression

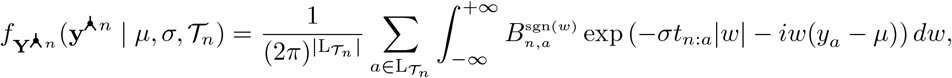

so that the recursion holds, which ends the proof.□

##### Proof of Proposition 1.

It is straightforward to prove by induction that for all nodes *n* of *𝒯* and all tips *a ∈ 𝒯*_*n*_, we have that 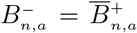. By setting 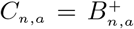 for all nodes *n* of *𝒯* and all tips *a ∈ 𝒯*_*n*_, Proposition S1 directly implies Proposition 1 of the main document.□

### B Supplementary Figures and Tables

#### B.1 Simulations: Model Selection

**Figure S1:**
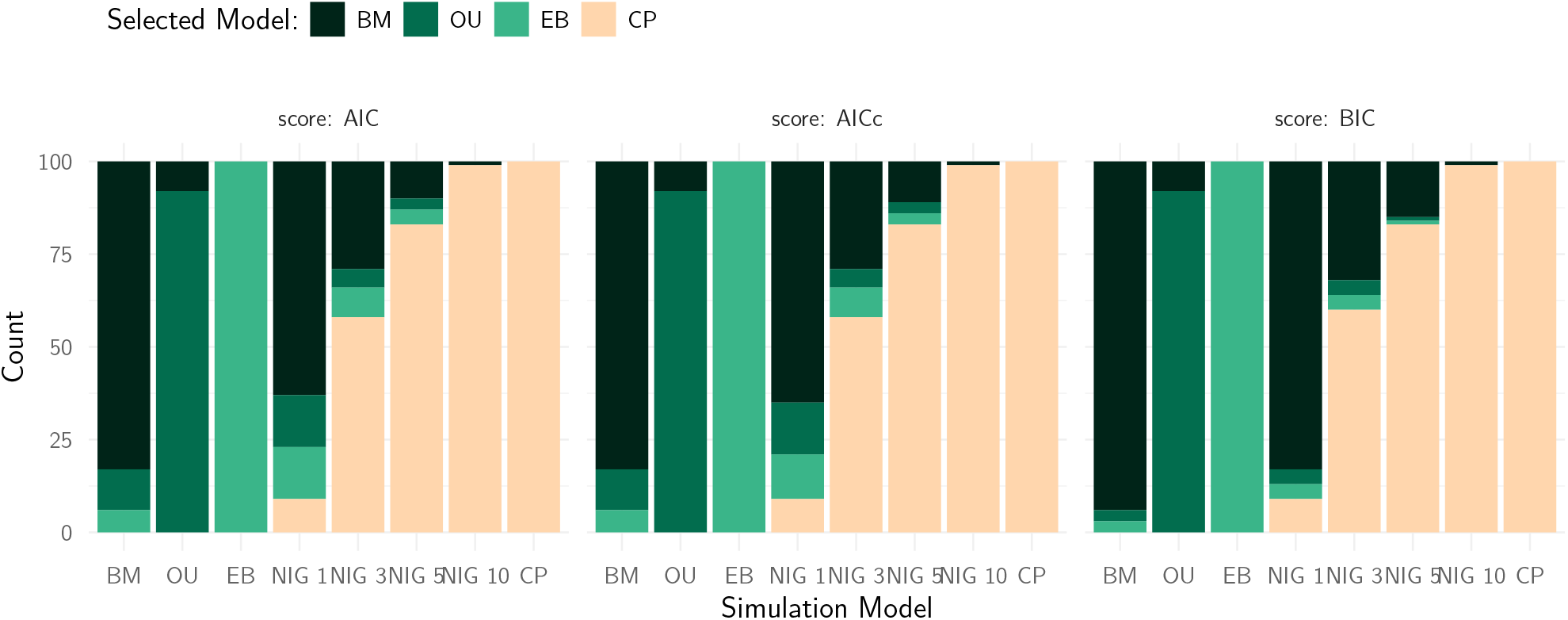
Model selection results using the AIC, AICc or BIC for 100 datasets simulated on the Lizard tree (Mahler et al., 2013), with three Gaussian processes (BM, OU, EB), the NIG process with various levels of excess kurtosis (from 1 to 10), and the Cauchy process. All simulation processes but the EB have the same MAD as the BM process with variances 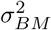 of 5, 10, 50 and 100. For each simulation model, the proportion of time each fitting model is selected is shown as a bar plot.

#### B.2 Simulations: Parameter Inference

**Figure S2:**
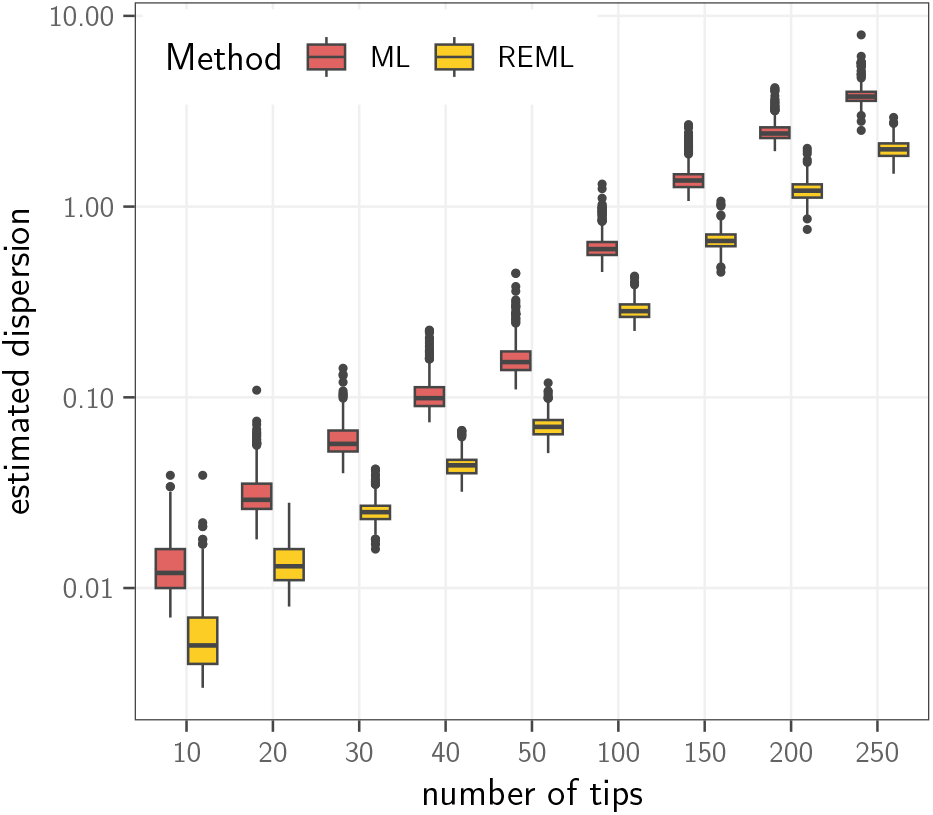
Time needed on a standard laptop computer (MacBookPro 13-inch, M2, 2022) for the fit of a CP on pure-birth trees of growing sizes (x axis), using the maximum likelihood with fixed root method (ML, dark red) or the restricted maximum likelihood (REML, light orange). Each boxplot is on 500 replicates.

#### B.3 Greater Antilles Lizards

**Figure S3:**
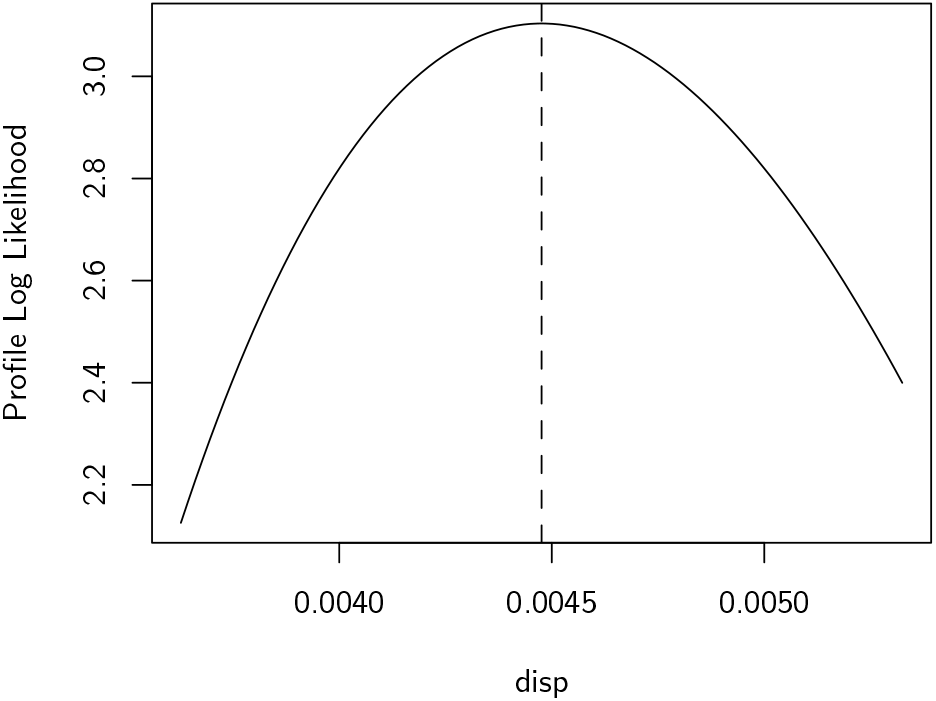
Profile log likelihood for the dispersion parameters of the svl trait, fitted with a CP using the REML. The likelihood curve is smooth and has a clear maximum, which is a sign that the numerical optimization and likelihood computations went well.

#### B.4 West Nile Virus

**Table S1:** Fits of Gaussian (BM, OU and EB) and CP models on the WNV dataset from Pybus et al. (2012), on the MMC tree for the latitude and longitude traits. The first two lines are the log likelihood and AIC scores for the ML fit. The parameters are *μ* for the ancestral root value, *σ* for the Gaussian standard error or Cauchy dispersion, and *θ* represents the selection strength *α* for the OU, and the rate *r* for the EB. The computation time for each fit on a standard laptop is given in seconds.

|  | latitude |  |  |  | longitude |  |  |  |
| --- | --- | --- | --- | --- | --- | --- | --- | --- |
|  | BM | OU | EB | CP | BM | OU | EB | CP |
| logLik | -285.68 | -285.68 | -277.31 | **-247.97** | -352.30 | -352.30 | -343.93 | **-308.45** |
| AIC | 575.37 | 577.37 | 560.62 | **499.95** | 708.60 | 710.60 | 693.85 | **620.90** |
| $\mu$ | 41.20 | 41.20 | 41.16 | **40.81** | -75.34 | -75.34 | -75.54 | **-74.16** |
| $\sigma$ | 2.80 | 2.80 | 7.78 | **0.40** | 5.31 | 5.31 | 15.67 | **0.73** |
| $\theta$ | – | 0.00 | -0.53 | – | – | 0.00 | -0.56 | – |
| time (s) | 0.00 | 0.02 | 0.01 | **2.04** | 0.00 | 0.01 | 0.02 | **2.00** |

**Figure S4:**
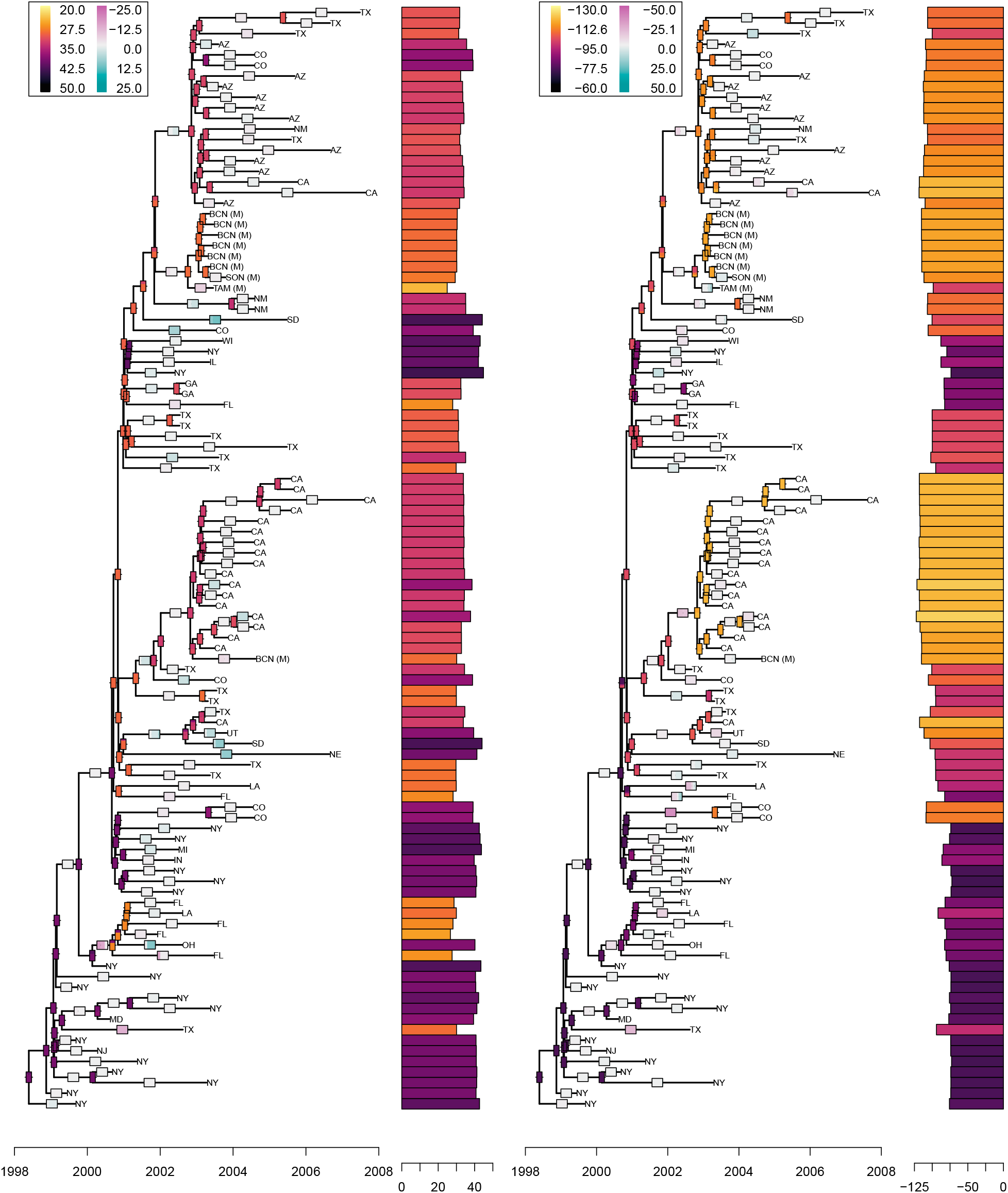
Ancestral trait reconstruction for the WNV dataset from Pybus et al. (2012), for the latitude and longitudes considered as two independent traits, using the Cauchy REML model fit. Measured traits are shown as a bar plot at the tips of the tree, with a color gradient encoding the trait value, from light yellow (small) to dark purple (large). Modes of the posterior trait reconstructions are showed at internal nodes, with a width proportional to the relative weight of the modes (nodes with only one color have a single mode), and the same color scale as tip values. Modes of the posterior branches increments are showed on the edges, using the same convention, but with a diverging color scale so that increments close to zero are white, positive increments are blue, and negative increments are red.

**Figure S5:**
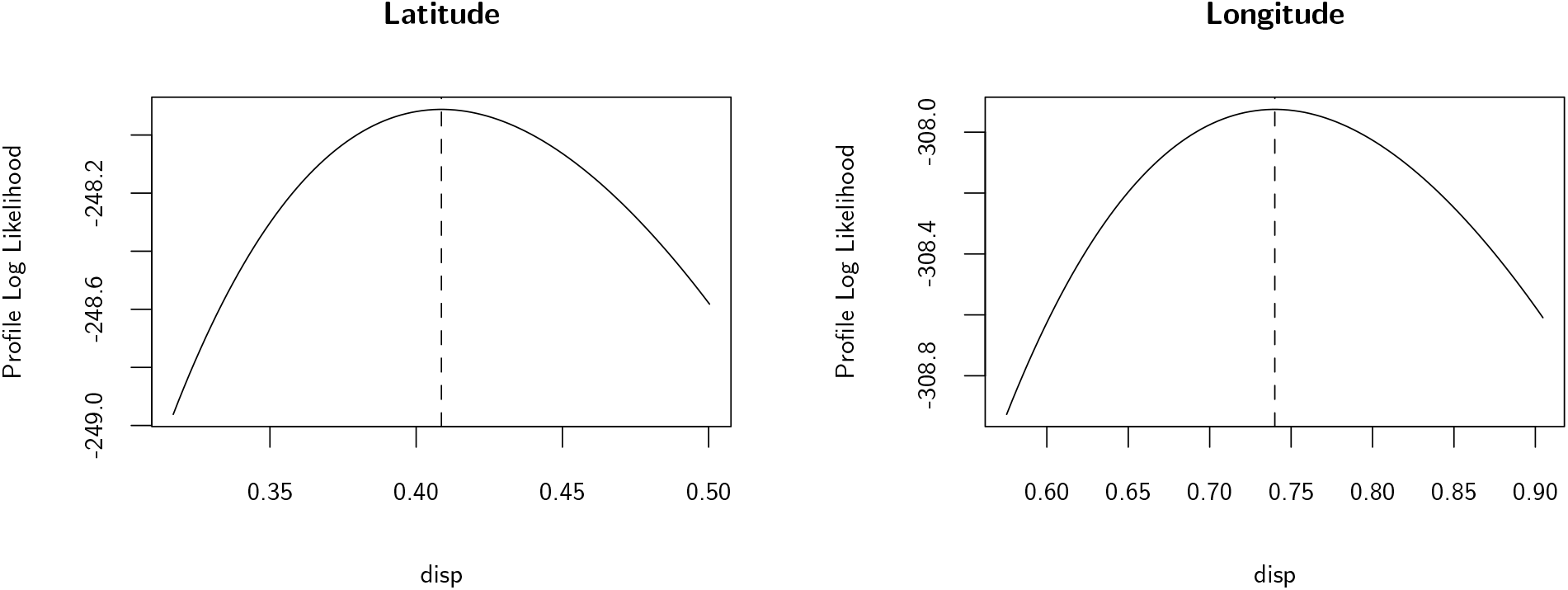
Profile log likelihood for the dispersion parameters of the latitude and longitude traits, both fitted separately with a CP using the REML. In both cases, the likelihood curve is smooth and has a clear maximum, which is a sign that the numerical optimizations and likelihood computations went well.

### C Model Selection with Higher Variance and Gaussian Noise

In this section, we use the same setting as in Section “Model Selection” of the main document, using the fixed empirical tree from the Lizard dataset (Mahler et al., 2013), normalized to unit height, and simulating tip trait datasets using the same evolutionary models, but we either raised the variance of the reference BM process, or added some independent Gaussian measurement error at the tips.

#### C.1 Higher Variance

In order to check how a higher variance or dispersion affects model selection, we simulated datasets with higher MAD values. We took the variance of the BM as the base parameter, that we varied from 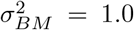 (as in the main text) to 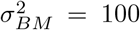. Using the same procedure as in Section “Model Selection” of the main document, we then selected the parameters of the other processes, so that they all had the same MAD values (except for the EB, see main text for more details).

We then applied the same inference and model selection procedure as in the main text for each newly generated tip trait datasets. Figure S6 displays the results obtained with reference BM variances 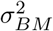 of 5, 10, 50 and 100 (associated MAD are, respectively, 1.51, 2.13, 4.77 and 6.74).

We can see that the graphics are very close one to another and to the results of the main text, with only slight stochastic variations. This suggests that variance or dispersion, i.e., evolution speed, does not influence the model selection accuracy. In particular, a BM with a high variance is not mistaken for a CP with small dispersion, and the jump signature of a CP cannot be captured by a BM (Fig. S6).

#### C.2 Gaussian Noise

In order to check to what extent an additional independent Gaussian noise affects model selection, we simulated datasets with the same parameters as the in Section “Model Selection” of the main document (i.e., with 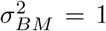), and then added an independent Gaussian error to the tip trait values resulting from each process. We varied the variance 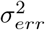 of the independent noise from 0.01 to 10.

We then applied the same inference and model selection procedure as in the main text for each newly generated tip trait datasets. Figure S7 displays the results obtained with reference Gaussian noise variances 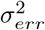 of 0.01, 0.1, 1 and 10.

As expected, adding a Gaussian error impairs the accuracy of the model selection, and high amounts of Gaussian noise tend to “cover” the signature of non-Gaussian processes. We however observe that almost 40% of the datasets simulated from a CP with dispersion 0.67 and a Gaussian error with variance 10 are still inferred as coming from a CP. This suggests that the jump signature of the Cauchy process is very strong, and is difficult to erase, even with high levels of Gaussian noises (Fig. S7).

As the *λ*-CP can accommodate for some independent (Cauchy) noise at the tip, we included it in the comparisons on this example (see the *Pagel’s lambda* paragraph of the Discussion of the main text). Here, it improved the results a little, raising the number of cases where the CP (with Cauchy errors) was correctly recovered over a Gaussian process (Fig. S8).

**Figure S6:**
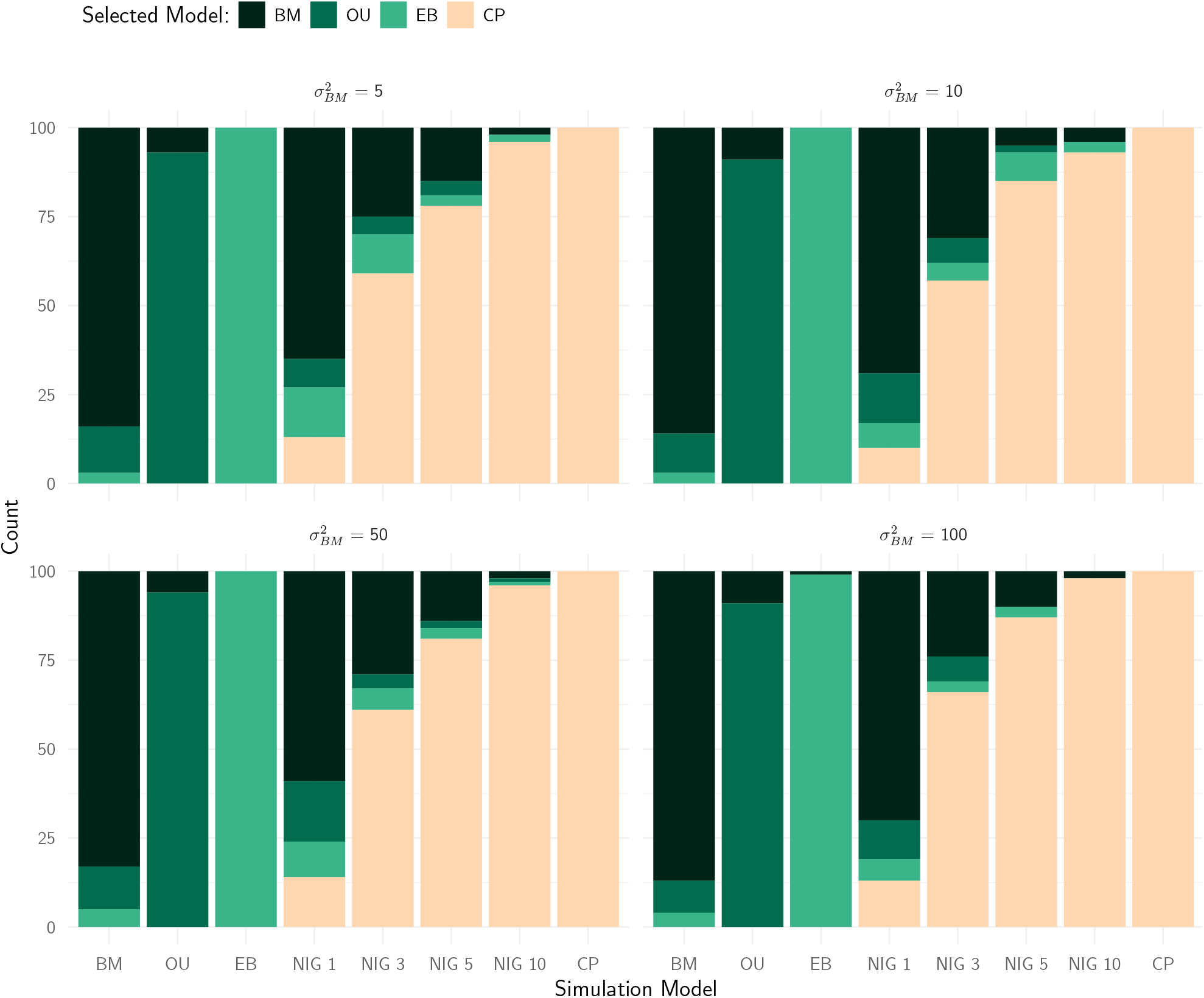
Model selection results using the AIC for 100 datasets simulated on the Lizard tree (Mahler et al., 2013), with three Gaussian processes (BM, OU, EB), the NIG process with various levels of excess kurtosis (from 1 to 10), and the Cauchy process. All simulation processes but the EB have the same MAD as the BM process with variances 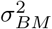 of 5, 10, 50 and 100. For each simulation model, the proportion of time each fitting model is selected is shown as a bar plot.

**Figure S7:**
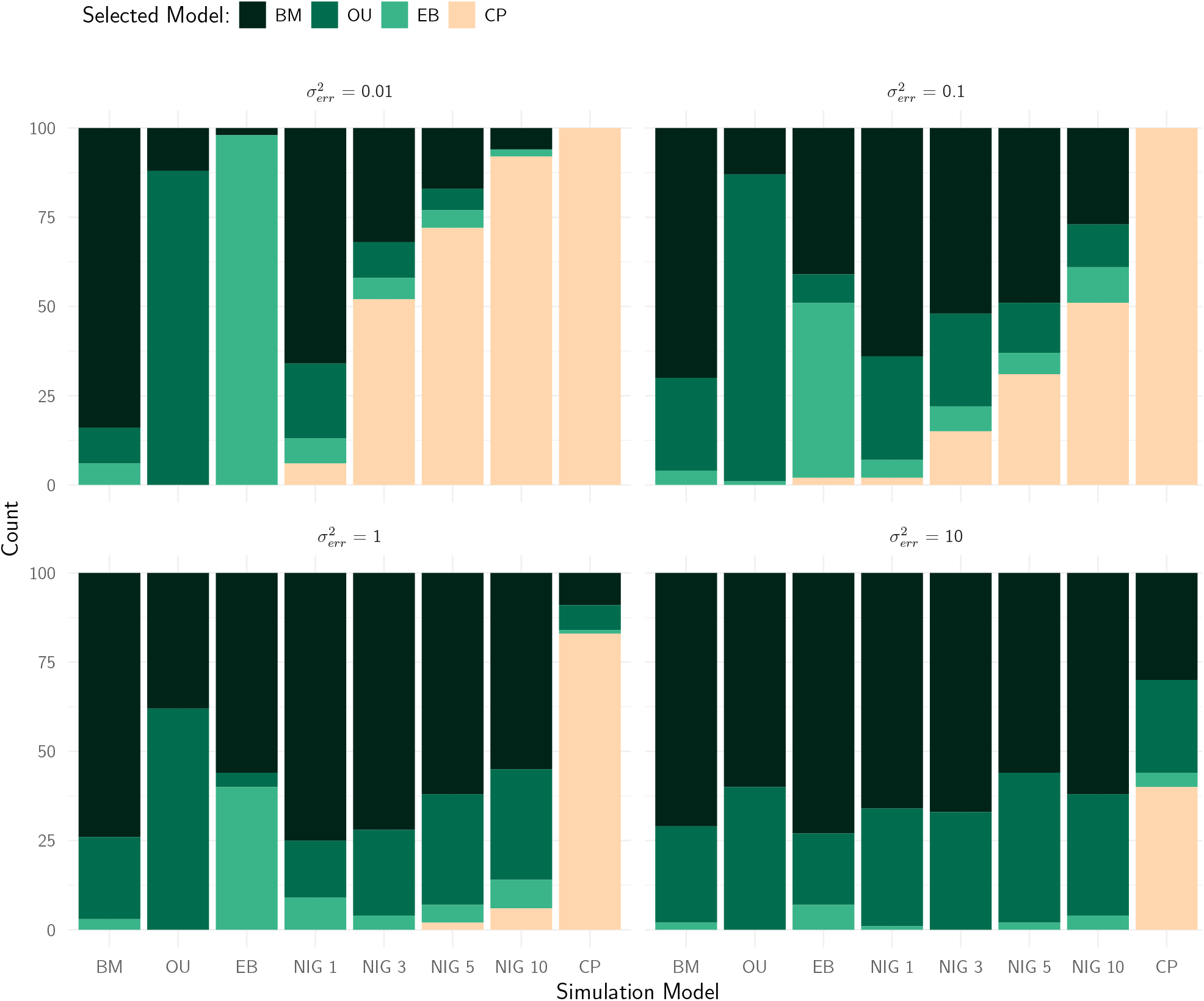
Model selection results using the AIC for 100 datasets simulated on the Lizard tree (Mahler et al., 2013), with three Gaussian processes (BM, OU, EB), the NIG process with various levels of excess kurtosis (from 1 to 10), and the Cauchy process. All simulation processes but the EB have the same MAD as the BM process with variance 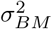 of 1. Independent Gaussian errors with variances 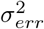 of 0.01, 0.1, 1 and 10 were added to all the tip values. For each simulation model, the proportion of time each fitting model is selected is shown as a bar plot.

**Figure S8:**
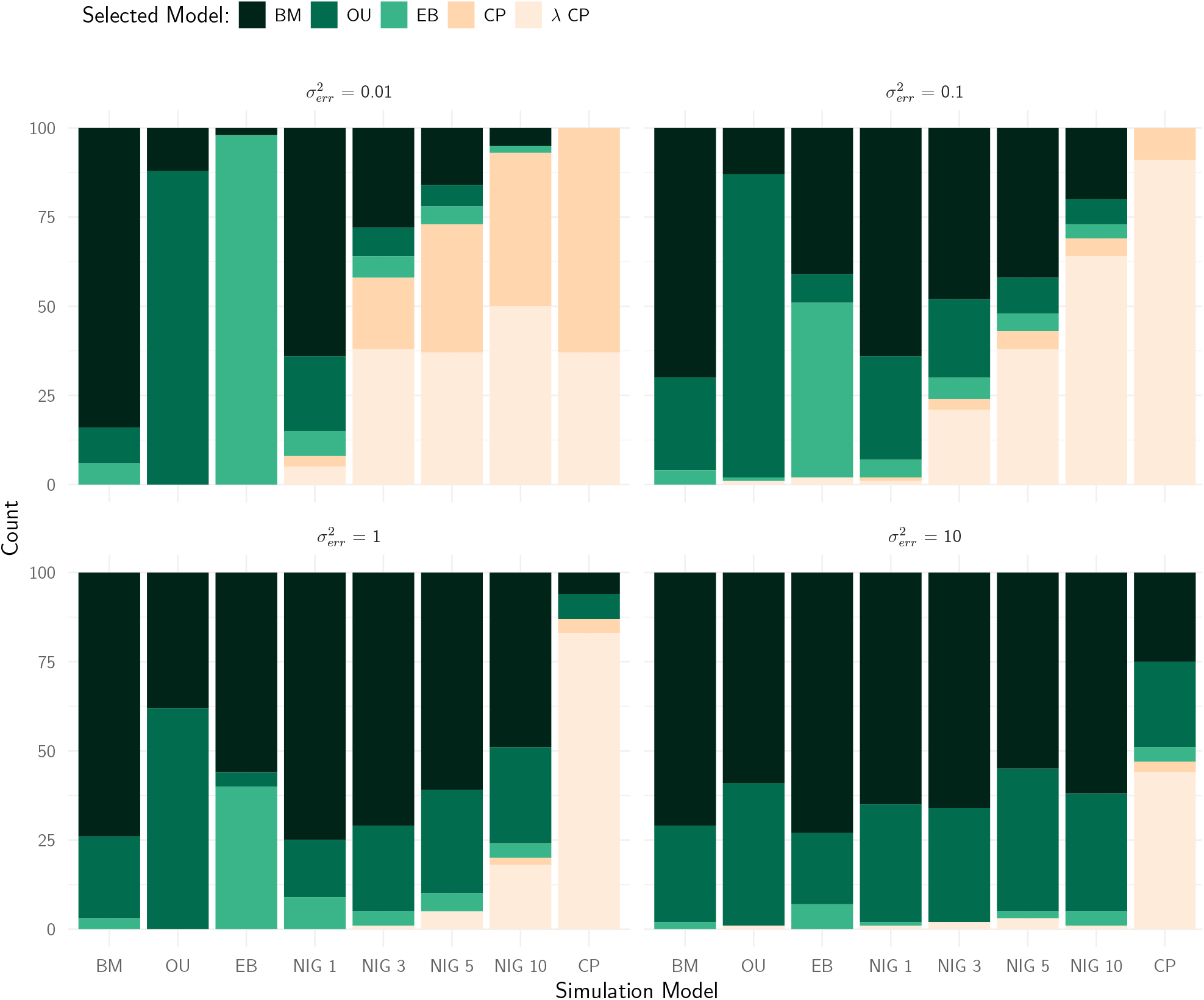
Same as Figure S7, but including the “*λ*-CP” in the model selection comparison. When the data is simulated by a CP with Gaussian noise of variance 10 (last column of the bottom right panel) The CP or *λ*-CP is selected 47 times out of 100, instead of only 40 times if the *λ*-CP is not considered.

## Notes

https://github.com/pbastide/cauchy_paper

https://github.com/gilles-didier/cauphy

### Competing Interest Statement

The authors have declared no competing interest.

### Summary of Updates

We corrected some typos. A new figure and additional analyses were added to the revised version of the manuscript.

